# Type I interferon responsive microglia shape cortical development and behavior

**DOI:** 10.1101/2021.04.29.441889

**Authors:** Caroline C. Escoubas, Leah C. Dorman, Phi T. Nguyen, Christian Lagares-Linares, Haruna Nakajo, Sarah R. Anderson, Beatriz Cuevas, Ilia D. Vainchtein, Nicholas J. Silva, Yinghong Xiao, Peter V. Lidsky, Ellen Y. Wang, Sunrae E. Taloma, Hiromi Nakao-Inoue, Bjoern Schwer, Raul Andino, Tomasz J. Nowakowski, Anna V. Molofsky

## Abstract

Microglia are brain resident phagocytes that can engulf synaptic components and extracellular matrix as well as whole neurons. However, whether there are unique molecular mechanisms that regulate these distinct phagocytic states is unknown. Here we define a molecularly distinct microglial subset whose function is to engulf neurons in the developing brain. We transcriptomically identified a cluster of Type I interferon (IFN-I) responsive microglia that expanded 20-fold in the postnatal day 5 somatosensory cortex after partial whisker deprivation, a stressor that accelerates neural circuit remodeling. *In situ*, IFN-I responsive microglia were highly phagocytic and actively engulfed whole neurons. Conditional deletion of IFN-I signaling (*Ifnar1^fl/fl^*) in microglia but not neurons resulted in dysmorphic microglia with stalled phagocytosis and an accumulation of neurons with double strand DNA breaks, a marker of cell stress. Conversely, exogenous IFN-I was sufficient to drive neuronal engulfment by microglia and restrict the accumulation of damaged neurons. IFN-I deficient mice had excess excitatory neurons in the developing somatosensory cortex as well as tactile hypersensitivity to whisker stimulation. These data define a molecular mechanism through which microglia engulf neurons during a critical window of brain development. More broadly, they reveal key homeostatic roles of a canonical antiviral signaling pathway in brain development.

## Introduction

Neural circuits undergo dynamic and experience dependent changes in circuit connectivity during brain development, and even subtle alterations in this process are associated with neurodevelopmental diseases^1–4^. Non-neuronal cells, including astrocytes and microglia, are essential to physiological neural circuit development and function ^5,6^. Microglia are the dominant immune cells in the brain parenchyma and key mediators of brain-immune communication, which is impacted in neurodevelopmental diseases including autism, epilepsy, and schizophrenia^7^. Microglia impact multiple aspects of neural circuit development, including synapse formation and elimination, elimination of whole cells, and modulation of neuronal activity^8,9^. Single-cell sequencing approaches have identified distinct subsets of microglia, many of which are conserved in both development and disease. However, in most cases, the functional implications of these molecular signatures is unknown.

One highly conserved microglial subset observed in aging, neurodegeneration, and other pathologies expresses a Type I-interferon (IFN-I) responsive signature^10^. This observation is puzzling, as the literature on IFN-I signaling is overwhelmingly focused on its roles in antiviral defenses^11^ and its function in homeostasis or sterile pathologies are not immediately obvious.

IFN-I cytokines are highly evolutionarily conserved, consisting of one IFN-beta transcript and multiple IFN-alphas, all of which signal through a receptor that includes the obligate IFNAR1 receptor subunit. IFN-I responsive microglia are an extremely small subset in most single-cell datasets, also raising questions as to its physiologic relevance. However, given that microglial states are highly dynamic to local factors, including cytokines, neuronal activity and damage-associated molecular patterns^5,12,13^, it is possible that low abundance subsets reflect transient but functionally critical cell states.

Here we found that a partial whisker lesion in early life led to a 20-fold expansion of a Type I interferon (IFN-I) responsive microglial subset (Interferon-responsive microglia, <u>IRMs</u>) in the developing somatosensory cortex. IRMs had a highly phagocytic morphology, with multiple projecting phagosomes per cell, and were often in the process of engulfing whole neurons *in situ*. Global or microglial-specific deletion of IFN-I sensing led to dysmorphic microglia with distended phagosomes, and an accumulation of cortical neurons harboring double strand DNA (dsDNA) breaks, a marker of cell stress or hyperexcitability. Conversely, exogenous IFN-β was sufficient to drive microglial engulfment of neurons and restrict the number of cortical neurons with dsDNA breaks. Loss of IFN-I sensing led to altered somatosensory neuron subtypes and tactile hypersensitivity in juvenile mice. Taken together, our data reveal a physiological role for IFN-I-driven microglial phagocytosis in brain development.

## Results

### A type I interferon-responsive microglial subset expands 20-fold during cortical remapping

To interrogate dynamic microglial responses we examined the rodent barrel cortex, a canonical and versatile model of developmental structural plasticity^14–18^. Lesions of the afferent whisker sensory neurons at postnatal day 2 (P2) lead to topographic remapping of the whisker representations in the contralateral cortex by P5^14,15,19^, during an active period of microglial proliferation and migration^20^. Importantly, because the whisker somatosensory circuit synapses in the brainstem and the thalamus en route to the cortex, whisker lesion promotes cortical remodeling without inducing an acute injury response. Complete whisker elimination leads to synaptic loss by P14 and has been used to discover microglial mechanisms of synapse elimination^21^. In this case, we established a subthreshold deprivation model, cauterizing 40% of whisker follicles at P2 (rows B and D) to promote circuit rearrangement but not synaptic loss (**Fig. 1A**).

**FIGURE 1:**
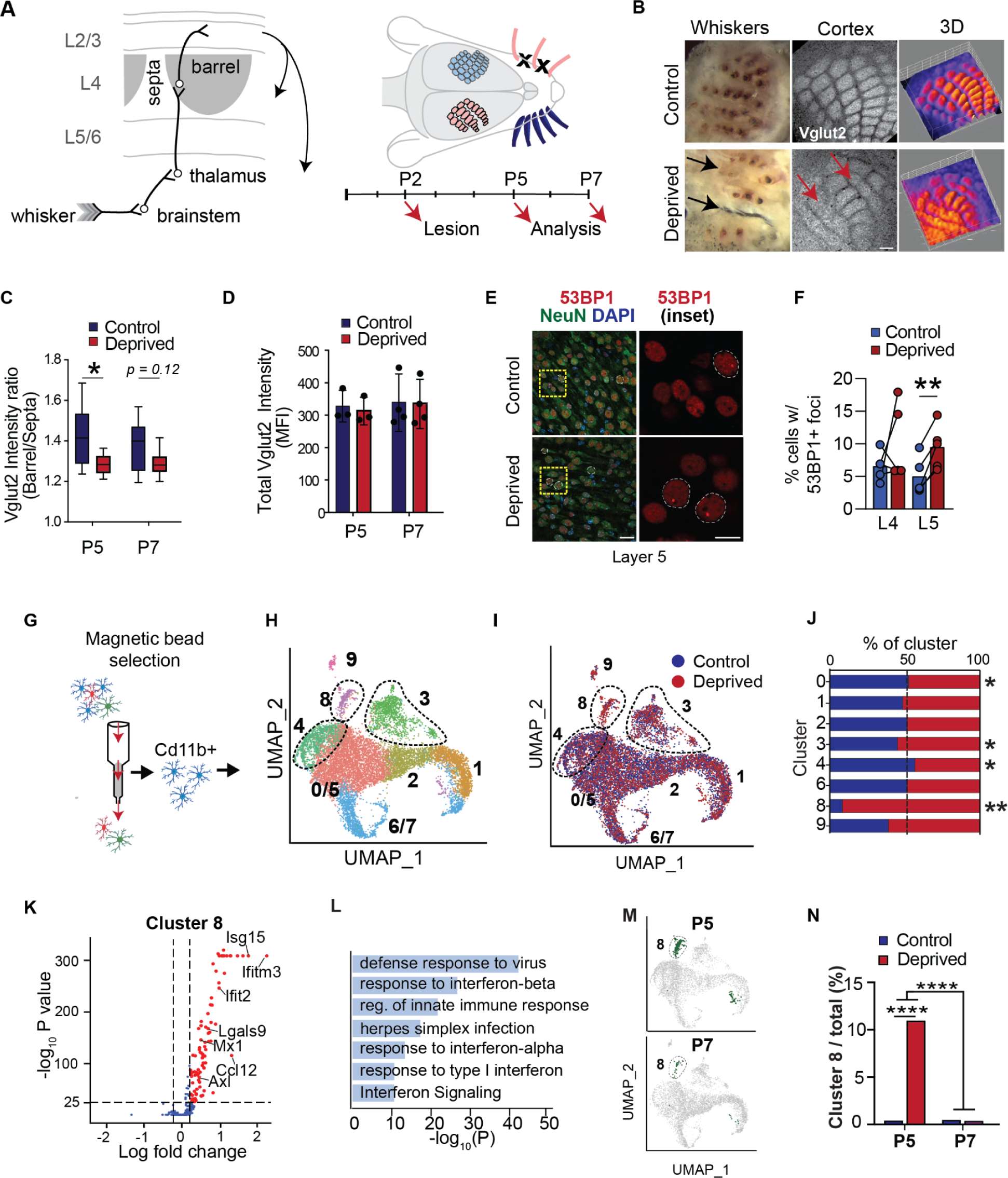
A type I interferon-responsive microglial subset expands 20-fold during cortical remapping. **(A)** Schematic of barrel cortex connectivity and experimental timeline **(B)** Representative images of control and whisker lesioned whisker pad (left), *en face* imaging of L4 somatosensory cortex and cortex topographical heat map derived from VGLUT2 intensity data (right). (scale bar = 100μm) **(C)** Quantification of barrel distinctness based on VGLUT2 intensity in barrels vs. septa in control and deprived hemispheres. Box and whisker plots show range (whiskers), median, and first and third quartiles (box). (2-4 barrel/septa pairs per condition per mouse; P5: n=3 mice, P7: n=4 mice, One-way ANOVA with Sidak post-hoc test, p=0.0380/0.1223) **(D)** VGLUT2 intensity averaged over the entire barrel field in the deprived and spared hemispheres. Error bars show mean±SD. (P5: n=3 mice, P7: n=4) **(E)** Representative images of 5BBP1+ foci in layer 5 of the barrel cortex. Inset of the right with 53BP1+ foci containing neurons circled in dotted white line. (scale bar = 20μm and scale bar inset = 10μm) **(F)** Quantification of 53BP1 foci+ neurons in layers4/5 of the barrel cortex in control vs. deprived hemispheres. Dots per mouse, line connect both hemispheres from same mouse. (n=5mice, one-way ANOVA with Sidak post-hoc test, p=0.5728/0.0055) **(G)** Schematic of MACS-isolation of microglia for single cell sequencing **(H)** UMAP plot showing independent clustering of 12,000 CD11b+ cells, pooling P5 and P7 timepoints from control and whisker deprived cortices, including microglia (Clusters 0-8) and macrophages (Cluster 9). Dotted lines highlight clusters for further analysis. Clusters 0/5 and 6/7 were the results of merging nearest neighbor clusters 0 with 5 as well as 6 with 7 due to low numbers of uniquely expressed genes, as shown in **Fig. S2E**. **(I)** Clusters colored by condition (control vs. whisker deprived) **(J)** Quantification of cluster composition by condition. X-axis represents percent of cells in each cluster from the control (blue) or deprived (red) hemispheres, normalized for total number of cells per sample. (Chi-square test with Bonferroni correction, *pAdj<0.01, **pAdj<10^-25^) **(K)** Volcano plot of genes differentially expressed in cluster 8 relative to all other clusters **(L)** GO terms upregulated in Cluster 8 (Metascape^81^) **(M)** UMAP plots showing cells separated by age (P5 vs. P7) with cluster 8 highlighted in green **(N)** Plot of cluster 8 abundance at each time point in control vs. deprived conditions (Chi-square test with Bonferroni correction, ****pAdj<10^-10^) *See also* ***Fig. S2***, **Supplementary Tables 2-3**

We first surveyed key aspects of circuit development in our partial deprivation modeL. As expected, neonatal whisker removal led to redistribution of the VGLUT2+ thalamocortical axonal boutons in cortical layer 4, whereby the deprived cortical rows (B and D) were smaller with indistinct barrel separation, compared to the contralateral (control) cortex (**Fig. 1B-C**, ^18^). However, mean VGLUT2 intensity across the barrel cortex was not significantly different, suggesting that overall thalamocortical synapse density was preserved (**Fig. 1D**). Since neuronal death has been reported in some variations of this model, we also examined cell death. We did not observe changes in markers of apoptosis or end stage DNA damage (**Fig. S1A-D**). However, we observed a significant increase in the proportion of neurons in layer 5 harboring dsDNA breaks using 53BP1(**Fig. 1E-F, Fig. S1E-F**), a protein involved in non homologous end joining that clusters at the site of dsDNA breaks. These lesions are frequently observed as an early stage marker of cell stress and damage, but have also been postulated to correlate with hyperexcitability^22–24^. Microglial numbers and IBA1 expression were unaltered, suggesting no overt microgliosis (**Fig. S1G-J**). Thus, this partial deprivation model leads to circuit remodeling without inducing overt synaptic loss or microglial reactivity.

To molecularly define the microglial response to topographic remapping we performed single cell RNA sequencing of microglia at P5 and P7 after whisker removal at P2. We purified microglia using mechanical dissociation followed by magnetic bead isolation, which is ideal for preserving *in vivo* microglia signatures (**Fig. 1G** ^25,26^). We recovered 12,330 cells from 10 mice after filtering, quality control, and *in silico* selection of brain myeloid cells using unbiased clustering and expression of marker genes (**Fig. 1H, Fig. S2A-D**). Two clusters with low cell numbers were manually combined with their nearest neighbors due to low numbers of uniquely enriched genes (clusters 0/5 and 6/7; **Fig. S2E**). We obtained eight clusters, four of which were altered by whisker deprivation **(Fig. 1I-J**, **Supplemental Tables 1-3**). Clusters 1, 2, and 6/7 clustered mainly by markers of the different stages of cell division (**Fig. S2G-H**). A small macrophage subset (cluster 9, *Pf4*, *Lyve1*) was not substantially changed by whisker deprivation. The remaining four non-proliferative clusters of microglia (0/5,3,4,8), all changed in relative abundance after cortical remodeling (**Fig. 1I-J, Fig. S2F,I-J**). These included a homeostatic cluster (*P2ry12, Ccr5*, cluster 0/5) and a cluster resembling ‘proliferative-region associated microglia’ and ‘damage-associated microglia’ (cluster 4, 38/42 ‘PAM’ genes upregulated; ^27–29^ 36/83 DAM genes upregulated (**Fig. S2K**)), both of which decreased after deprivation. A cluster containing canonical neuronal genes (cluster 3, *Rbfox3, Grin1)* and was modestly increased after whisker deprivation.

The most striking difference between control and whisker-deprived conditions was the emergence of a microglial subpopulation enriched in type I interferon (IFN-I) response genes (cluster 8; **Fig. 1I-L, Fig. S2M-N**). This cluster was located between microglia and macrophages in UMAP space but showed microglia-like expression of *Tmem119*, *Tgfbr1*, and *Hexb* (**Fig. S2L**). IFN-I signaling is a highly evolutionarily conserved antiviral response, but its functions in the developing brain are unknown^11^. Differential gene expression and Gene Ontology (GO) analysis on cluster 8 revealed a robust interferon response signature (*Ifitm3*, *Mx1*, *Ifit3, Isg15*, *Irf7,* and *Stat1)* and GO terms “Response to virus” (GO:0009615), “Response to interferon-alpha” (GO:0035455), and “Response to interferon-beta” (GO:0035458) (**Fig. 1K-L**). The interferon-responsive cluster 8 was enriched 20-fold in P5 deprived vs. control cortices (0.5% in control vs 11% in deprived) but was indistinguishable from control by P7 (0.6% vs 0.4%) (**Fig. 1K-L, Fig. S2M-P**). These data reveal a microglial subset that is rare in the typically developing cortex but expands markedly during a restricted phase of topographic remapping.

### Interferon-responsive microglia are conserved across pathologies

To determine if this IFN-I-responsive microglial subset correlated with previously described IRMs in pathological settings we examined murine microglial bulk transcriptomic datasets from a variety of physiological and pathological conditions. We analyzed the gene signature upregulated in Cluster 8 (**Fig. 1K**) within each of the published datasets and highlighted *Ifitm3* and *Mx1*, two of the classically upregulated genes in IFN-I signaling (**Fig. 2A, Supplemental Table 4**, ^30^). As expected, the IFN-I signature was most enriched after viral infection (lymphocytic choriomeningitis, LCMV). However, we also detected prominent induction of an interferon response in various sterile pathologies, including mouse models of brain tumors and of Alzheimer’s disease (AD). We observed similar results when comparing the IFN-I gene signature with recently published single cell datasets including AD models, glioblastoma models, Sars-CoV-2 infection, demyelinating injury (LPC, Cuprizone treatment), aging, and middle cerebral artery occlusion (**Fig. 2B-C**, **Supplemental Table 5**, ^27, 30–37)^.

**Figure 2:**
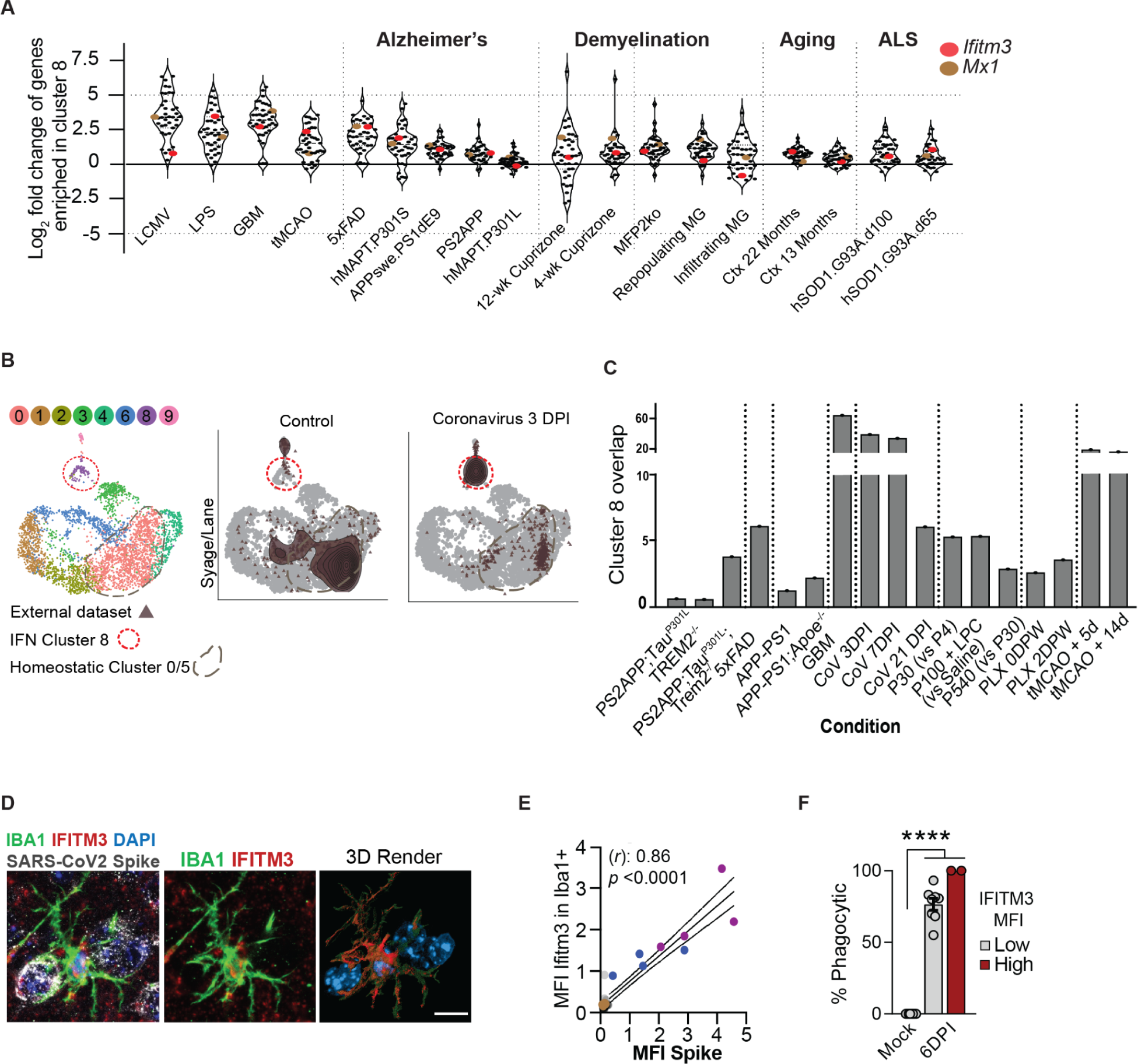
Interferon-responsive microglia are conserved across pathologies. **(A)** Expression of 38 ‘Cluster 8’ marker genes (eigengene) across a set of published microglia sequencing data (from bulk sorted CD11b+ cells^30^. Dots represent individual genes, highlighting *Ifitm3* (red) and *Mx1* (orange). Y-axis represents the log fold change relative to its own control. See Table S4 for details of each sample set and experimental condition. **(B)** Our microglial P5/P7 dataset was re-clustered to create a reference PCA and UMAP to which external datasets (brown triangles) were aligned. Red circle=Interferon-reponsive microglia (cluster 8), grey circle=homeostatic microglia. Representative examples shown are of a Sars-CoV2 dataset (see Table S5 for details). **(C)** Bar plot showing enrichment of Cluster 8-like cells in various microglial single-cell sequencing datasets from different disease states and ages relative to each study’s control. See Table S5 for all dataset descriptions and references ^27,31–37^ **(D)** Representative image and 3D reconstruction of an IFITM3+ microglia engulfing a SARS-CoV-2 infected cell (scale bar = 10μm) **(E)** Correlation of SARS-CoV-2 Spike protein with microglial IFITM3 mean fluorescence intensity per image. Each color represents an individual mouse. (n = 3 mice at 6 days post infection, r Spearman correlation coefficient = 0.86, p<0.0001) **(F)** Percent phagocytic microglia in cortices of mock-infected vs. SARS-CoV-2 infected mice at 6DPI, binned by IFITM3 expression per cell (see Methods for details). Dots per image. Kruskal-Wallis test, p<0.0001, n = 2 mice per group.

To further validate these findings we performed immunostaining of tissue sections for IFITM3, the top upregulated gene in IFN-I responsive cluster 8 (*Ifitm3,* **Fig. 1K**), in select models of pathology. This IFN-α/β - stimulated gene encodes a transmembrane protein known to limit viral replication, cancer progression, and amyloid beta plaque deposition^38–40^. Given its major public health interest, known involvement of IFITM3^41,42^, and recent reports of potential neurologic sequelae^43,44^, we examined the presence of IFITM3+ microglia in mice with neurotropic SARS-CoV-2 infection (K18-ACE2 transgenic mice^45–47^; **Fig. 2D, Fig S3A-D**). We observed high levels of IFITM3 in most microglia in brains with active viral replication, (**Fig. S3E**; ^48^). Microglia expressing higher levels of IFITM3 clustered around infected cells (**Fig. 2D**) and were more likely to have an amoeboid morphology (**Fig. S3F**). In summary, an IFN-I responsive microglial signature that emerges transiently during cortical development is conserved across multiple pathologic settings.

### IFN-I responsive microglia engulf neurons during cortical development

IFN-I responsive microglia (IRMs) are a molecularly conserved subset of microglia whose function is unknown. Using IFITM3 as a marker for this population (**Fig. 3A**), we examined their presence *in vivo* during development. We identified a rare population of IFITM3+ microglia in deeper layers of the developing barrel cortex (layers 4/5) that were increased up to 8-fold during topographic remapping, as quantified *in situ* and by flow cytometry (**Fig. 3B-C**, **Fig. S4A-B**). We confirmed IFITM3 is a bona-fide marker of our transcriptomically defined IRM population using several independent cluster markers. The canonical interferon stimulated transcription factor *Mx1* was also enriched after whisker deprivation *in vivo* as assessed by flow cytometry using an Mx1^GFP^ reporter^49^ (**Fig. S4A-B**). An alternate cluster 8 marker, BST2 (a.k.a. Tetherin^49,50^), was also increased after whisker deprivation and was expressed in most IFITM3+ cells, but was less specific than IFITM3 (**Fig. S4C-F**). Based on its similarities in developmental abundance, induction with barrel remodeling, location near barrels, and high concordance with two other markers that define our single cell cluster, we conclude that IFITM3 is a sensitive and specific marker that can identify the IRMs *in situ* during development.

**Figure 3:**
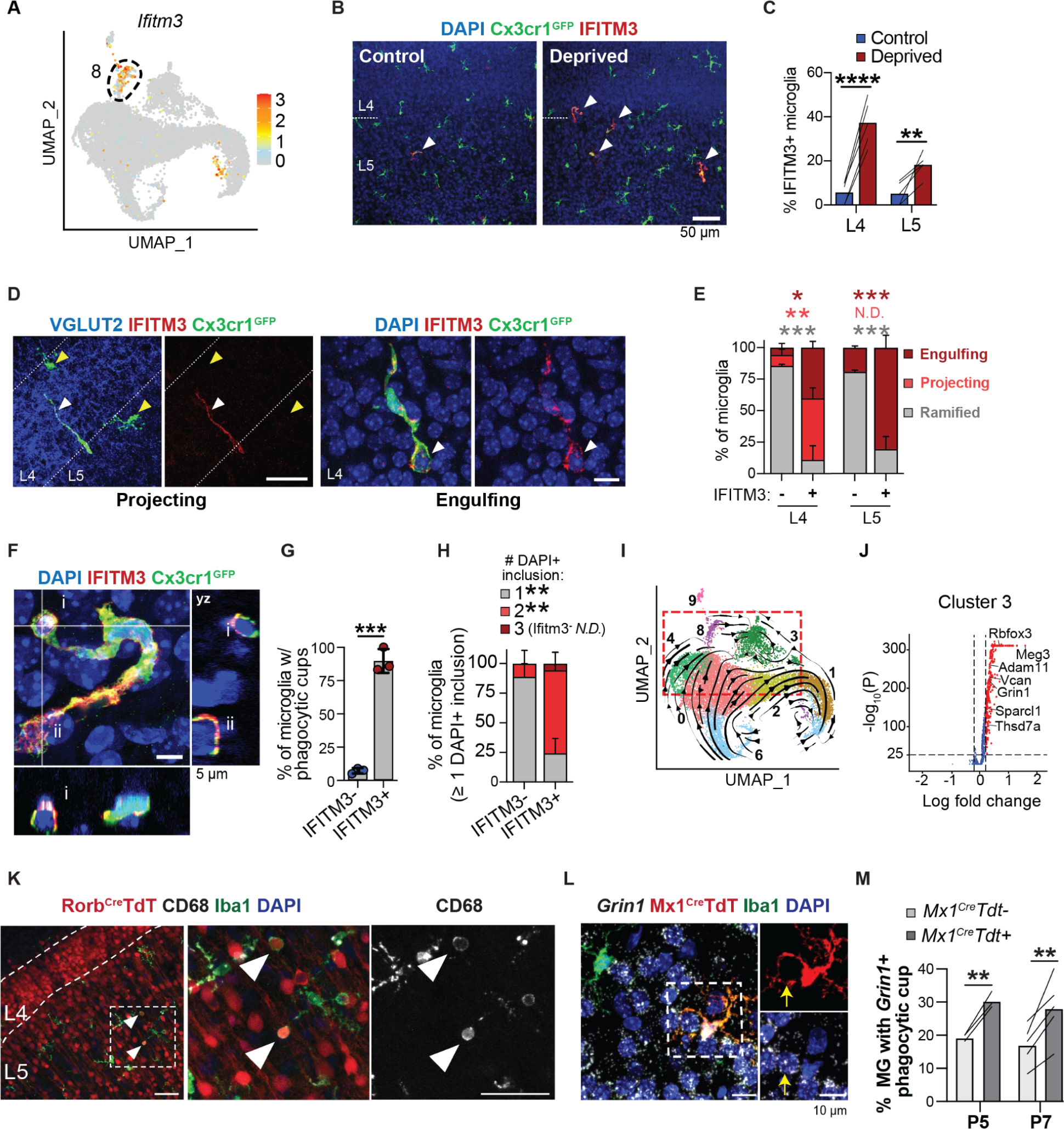
IFN-I responsive microglia engulf neurons during cortical development. **(A)** UMAP feature plot showing normalized expression of cluster 8 marker gene *Ifitm3* **(B)** Representative image of IFITM3+ Cx3cr1^GFP+^ microglia (arrowheads) in control or whisker deprived barrel cortex at P5 (scale bar = 50μm) **(C)** Percent IFITM3+ microglia from layers 4/5. Lines connect control and deprived hemispheres from the same mouse (n=5 mice, 2-way RM ANOVA with Sidak post-hoc test, p= <0.0001/0.0048) **(D)** Representative image of ‘projecting’ and ‘engulfing’ IFITM3+ microglia in P5 deprived cortex (scale = 20 μm (left), 5 μm (right)) **(E)** Quantification of microglial morphological subtypes described in Fig. 3D (n = 4 mice, 2-way RM ANOVA with Sidak post-hoc test, L4: p=0.0173/0.0086/0.0003 and L5: p=0.0005/0.9999/0.0005) **(F)** Confocal image and orthogonal views of an IFITM3+ microglia containing multiple phagocytic cups (i, ii show nuclei-containing phagocytic cups that are distinct from the microglia nucleus) (scale bar = 5μm) **(G)** Percent of IFITM3-and IFITM3+ microglia containing phagocytic cups (n = 3 mice, Welch’s t-test, p=0.0001) **(H)** Percent of microglia containing 1, 2 or 3 DAPI+ phagocytic compartments (inclusions), among microglia with at least one DAPI+ phagocytic compartment (n = 4 mice, 2-way RM ANOVA with Sidak post-hoc test, p=0.0029/0.0057/0.9772) **(I)** RNA velocity analysis showing predicted future cell state, colored by cluster (scVelo^54^) (Supplemental table 6) **(J)** Volcano plot of differentially expressed genes in Cluster 3 **(K)** Representative image of IBA1+ microglia engulfing RorbCre;TdT+ neurons in barrel cortex. Inset: layer 5. Arrowheads: Rorb^cre^TdT+ and CD68+ phagolysosomes (scale bar =50 μm) **(L)** Representative image of a tdTomato+ (*Mx1-Cre;Rosa26-LSL-TdT*) IBA1+ microglial phagocytic cup around a *Grin1+* nucleus (inset, arrow) (scale bar =10μm) **(M)** Quantification of percent tdTomato+ or negative microglia forming phagocytic cups around *Grin1+* nuclei in layers 4/5. (n=3 mice at P5 and 5 mice at P7, 2-way RM ANOVA with Sidak post-hoc test, p=0.0099/0.0029)

In contrast to the typical ramified morphology of most cortical microglia, IRMs were strikingly morphologically distinct. A subtype in L4 (‘projecting’) displayed an elongated morphology that appeared to project towards L5. A second subset enriched in L5 contained prominent IFITM3+ phagocytic cups enveloping DAPI+ nuclei (‘engulfing,’ **Fig. 3D-E**). These phagocytic IRMs often contained multiple DAPI+ inclusions, which appeared only rarely in IFITM3-microglia (**Fig. 3F-H**). IRMs had increased expression of the lysosomal marker CD68, consistent with increased phagolysosomal activity^51^ (**Fig. S4G-H**). IRMs were more likely to contain phagocytic cups than IFITM3-microglia (90% vs. 7%; **Fig. 3F-G**), and often had multiple phagosomes per microglia (**Fig. 3H**). IRMs were also more likely to contain both early and late stages of phagocytosis in the same cell (**Fig. S4I-J**). Phagocytosis proceeds in stages which include enveloping of extracellular material by the phagocytic cup, packaging into a phagosome, fusion to mature lysosomes to form a phagolysosome, and resolution of the phagolysosome by fragmentation^52,53^. Our data suggests that multiple stages of phagocytosis could be impacted by IFN-I signaling. For example, IFITM3 was correlated with early stages of engulfment (**Fig. S4K-L**) while BST2 was enriched in an intracellular perinuclear compartment (**Fig. S4D**). Taken together, these data suggest that IFN-I amplifies a microglial whole cell engulfment response enriched in cortical L5.

We next sought to investigate which cortical cell types were engulfed by IRMs. RNA velocity analysis of our single-cell dataset, which uses the ratio of unspliced pre-mRNA to spliced mRNA to infer trajectories between neighboring clusters^54^ predicted a trajectory from IRMs towards microglial cluster 3, which was enriched for neuronal mRNAs (*Rbfox3, Grin1, Gria1*; **Fig. 3I-J, Supplemental Table 6**). These neuronal mRNAs were enriched for intronic sequences associated with unspliced transcripts (**Fig. S4M-N**) suggesting that they were likely derived from engulfment of nuclear material rather than microglial expression of these genes.

To test if microglia could engulf neurons *in situ* we examined a *Rorb^cre^;R26R-TdT* line labeling most excitatory neurons in L4/5/6 (^55^; Jax 023526). Nine percent of L4/5 microglia had TdT-positive CD68-containing phagolysosomes at P5, suggesting that they had recently engulfed neuronal soma (**Fig. 3K, Fig. S4R**). In contrast, we observed no TdT inside microglia in an astrocyte reporter line at this age (*Aldh1l1^TdT^*) (**Fig. S4Q-R**). As independent validation, we performed *in situ* hybridization for *Rbfox1* and *Grin1*, two of the most abundant neuronal transcripts in cluster 3. Around 80% of phagocytic cup-containing microglia surrounded *Grin1+* neuronal soma at P5, declining to 20% at P7 (**Fig. S4O-P**). Finally, we directly examined whether an IFN-I responsive state in microglia precedes neuronal engulfment by fate mapping IRMs using an *Mx1^Cre^;R26R-TdT* reporter (Jax 003556 ^56^). As expected, Mx1^cre^TdT+ microglia increased after whisker deprivation; **Fig. S4S-T**). These Mx1^cre^TdT+ microglia were significantly more likely to contain phagocytic cups with the neuronal *Grin1* mRNA than TdT-cells (**Fig. 3L-M**). Given the increase in DNA-damaged neurons observed after whisker deprivation, we also wondered whether IRMs displayed a preference for damaged over intact neurons. Using the dsDNA break marker γH2AX (**Fig. S1E-F**), we found that IFITM3+ cells preferentially contacted neurons with dsDNA breaks^57^ (**Fig. S4U-V**). Together these data suggest that IFN-I responsive microglia in the barrel cortex are engulfing neurons *in vivo*.

### IFN-I signaling promotes microglial phagocytosis and restricts accumulation of DNA-damaged neurons

To determine whether IFN-I signaling was required for microglial phagocytosis and neuronal engulfment we examined mice lacking the obligate IFN-I receptor *Ifnar1* (Jax 028288; **Fig. 4A**). These mice completely lacked IRMs, consistent with a canonical IFN-I response (**Fig. 4B**). Microglia from *Ifnar1^-/-^* mice were markedly dysmorphic. They frequently contained multiple phagocytic compartments per cell and enlarged phagocytic compartments with thin walls enclosing diffuse DAPI+ nuclear material (**Fig. 4C-D, Fig. S4A-B**, **Supplemental Movie 1**). In contrast, wild-type microglia contained smaller phagocytic compartments with condensed DAPI signal completely enveloped by the cell membrane. This phenotype was reminiscent of ‘bubble’ microglia observed in zebrafish with deficits in phago-lysosomal fusion^58^. We defined bubble microglia in our model as those containing phagosomes larger than the cell nucleus (**Fig. 4E-F**). Bubble microglia were rare in control mice, but represented up to 50% of barrel cortex microglia from *Ifnar1^-/-^* mice (**Fig. 4F-G**). This phenotype was developmentally restricted to an early postnatal period (P5-P7) and was no longer detectable by P15 (**Fig. 4H**, **Fig. S5F**). Dysmorphic microglia were also increased in other regions of the *Ifnar1^-/-^* brain, including the corpus callosum and thalamus, and there were no changes in overall microglial density in any brain region (**Fig. S5C-E**). Interestingly, we also found a higher percentage of cells with dsDNA breaks in the barrel cortex of *Ifnar1^-/-^* mice than in wild type mice at P5 (**Fig. 4I-J**), indicating an accumulation of damaged neurons in IFN-I deficient brains. Similar to the bubble microglia phenotype, this increase in dsDNA breaks was restricted to this early postnatal window and no longer detectable in either genotype by P15 (**Fig. 4K**).

**Figure 4:**
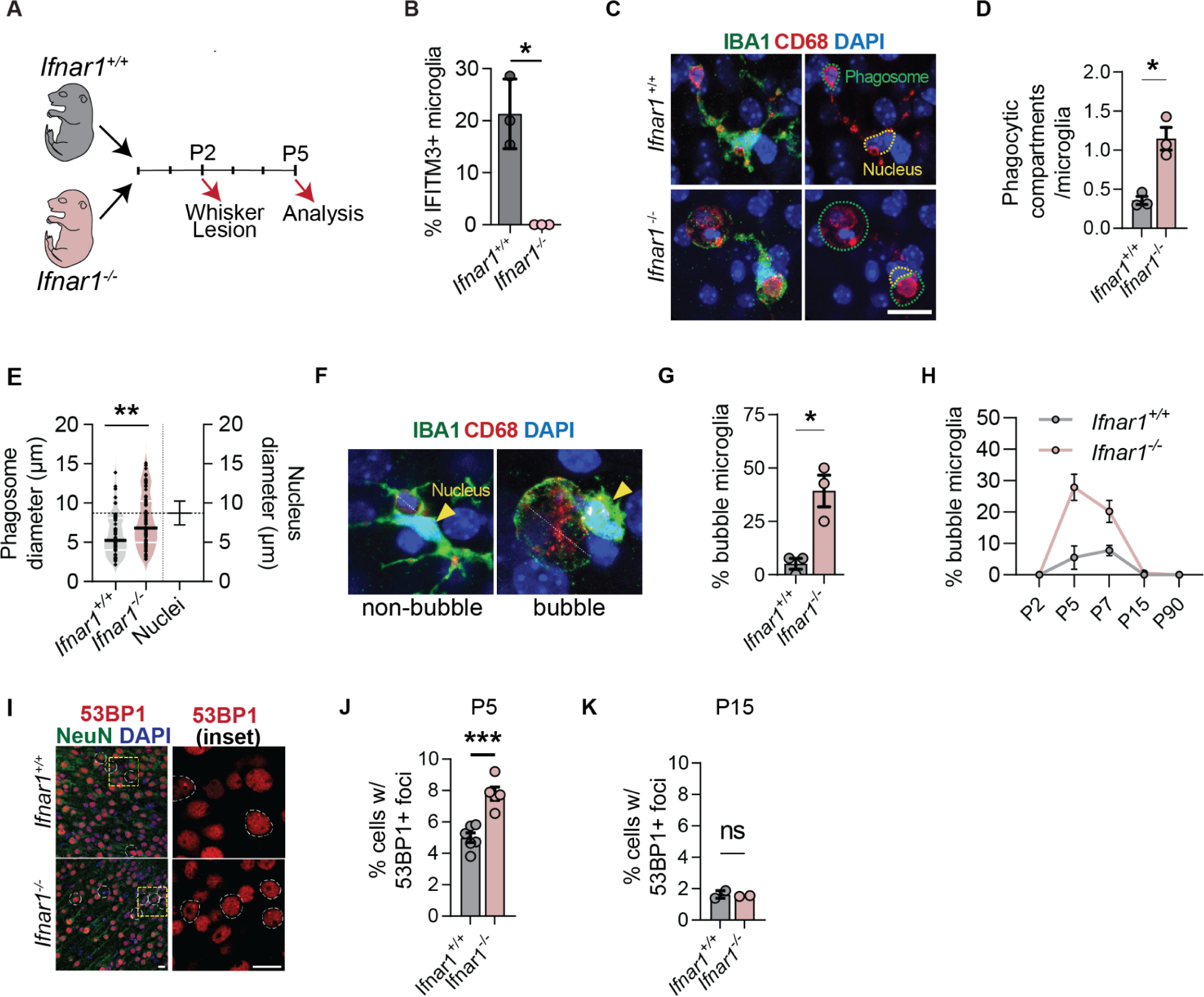
IFN-I signaling promotes microglial phagocytosis and restricts accumulation of DNA-damaged neurons. **(A)** Diagram of experimental design comparing control *(Ifnar1^+/+^)* vs. IFN-I insensitive *(Ifnar1^-/-^)* mice. **(B)** IFITM3+ microglia in deprived hemispheres L4/5 barrel cortex from *Ifnar1^+/+^* and *Ifnar1^-/-^*. Dots per mouse (Welch’s t-test, p=0.0313, n=3 per group) **(C)** Representative image of microglia from *Ifnar1^+/+^* and *Ifnar1^-/-^* mice. Yellow: microglia nucleus, green: phagocytic compartment (scale bar = 15µm) **(D)** Phagocytic compartments per microglia in L4/5 barrel cortex from *Ifnar1^+/+^* and *Ifnar1^-/-^* mice. Dots per mouse. Welch’s t-test, p=0.0228, n=3 mice per group. **(E)** Largest phagosome diameter per microglia in µm in L4/5 barrel cortex from *Ifnar1^+/+^* and *Ifnar1^-/-^* mice. Horizontal dotted line shows mean microglial nucleus size. Welch’s t-test, p=0.0011, n=57-76 microglia from 3 mice per group **(F)** Representative images of a non-bubble microglia containing a phagosome and a bubble microglia with an enlarged phagosome. Yellow arrowhead: microglia nucleus, white dashed line: phagosome diameter (scale bar = 10μm) **(G)** Percent bubble morphology microglia in L4/5 barrel cortex from *Ifnar1^+/+^* and *Ifnar1^-/-^* mice. Dots per mouse. Welch’s t-test, p=0.0332, n=3 mice per group **(H)** Developmental time-course of bubble morphology microglia in L4/5 barrel cortex from *Ifnar1^+/+^* and *Ifnar1^-/-^* mice. n= 2-3 mice per time point **(I)** 53BP1+ neurons in barrel cortices of *Ifnar1^+/+^* and *Ifnar1^-/-^* mice. Yellow dotted lines outline nuclei with 53BP1+ foci, square inset highlights 53BP1 staining. (scale bar = 10μm). **(J)** Percent of all cells in layers 4/5 containing foci in *Ifnar1^+/+^* and *Ifnar1^-/-^* mice at P5 (n= 5-6 mice, Welch’s t-test, p=0.0008). **(K)** Percent of all cells in layers 4/5 containing foci in *Ifnar1^+/+^* and *Ifnar1^-/-^* mice at P15 (n= 2 mice, Welch’s t-test, p=0.7446).

### IFN-I signals directly to microglia to promote neuronal engulfment and restrict accumulation of dsDNA break neurons

To determine potential cellular targets of IFN-I signaling we performed *in situ* hybridization for *Ifnar1* (**Fig. 5A**). We detected *Ifnar1* mRNA ubiquitously across cell types including microglia, neurons, astrocytes, and endothelial cells (**Fig. 5B-C**). Given both neuronal and microglial phenotypes, we tested which cell type was mediating these effects by conditional deletion of *Ifnar1 (Ifnar1^fl/fl^*; Jax #028256) from both cell types. We found that loss of IFN-I sensing by microglia phenocopied the bubble microglia phenotype observed in the knockout model (**Fig. 5D-E**; *Ifnar1^fl/fl^:Cx3cr1^Cre^*; MMRRC036395, vs. *Ifnar1^fl/fl^* control). These mice also phenocopied the increase of neurons with dsDNA breaks in layers 4/5 of the barrel cortex (**Fig. 5F**). In contrast, mice lacking IFN-I sensing by neurons had no increase in bubble microglia or neurons with dsDNA breaks (**Fig. 5G-H**; *Ifnar1^fl/fl^:Syn1^Cre^*; Jax003966). Taken together, these data suggest that direct IFN-I signaling to microglia is required for efficient phagocytosis and to limit damaged neurons in the developing cortex.

**Figure 5:**
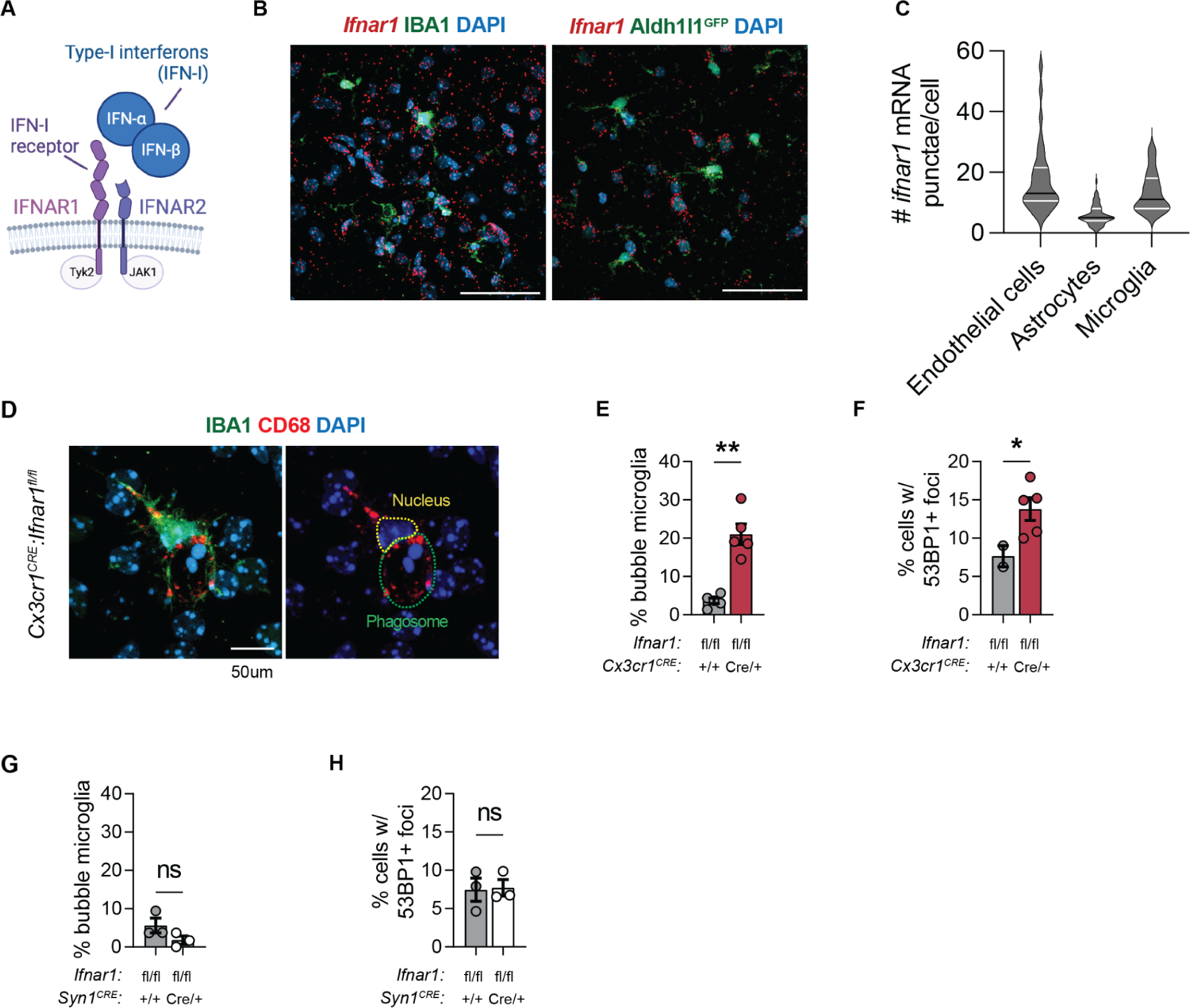
IFN-I signals directly to microglia to promote neuronal engulfment and restrict accumulation of dsDNA break neurons. **(A)** Schematic of Type I interferon receptor. **(B)** Representative confocal image of *Ifnar1* transcript co-stained with IBA1 or Aldh1l1^GFP^ at P5 in the somatosensory cortex (scale bar = 50μm) **(C)** Quantification of the number of punctae in nuclei of microglia (IBA1+), astrocytes (Aldh1l1^GFP+^) and endothelial cells (based on elongated morphology) **(D)** Representative confocal image of IBA1+ microglia in L4/5 barrel cortex from *Cx3cr1^CRE^:Ifnar1^flox/flox^* showing bubble phenotype with enlarged DAPI+ containing phagosome, Yellow: microglia nucleus, green: phagocytic compartment. (scale bar = 50μm) **(E)** Percent bubble morphology microglia in L4/5 barrel cortex from *Cx3cr1^CRE^:Ifnar1^flox/flox^* mice. Dots per mouse. (n=5mice, Welch’s t-test, p=0.002) **(F)** Percent bubble morphology microglia in L4/5 barrel cortex from *Syn1^CRE^:Ifnar1^flox/flox^* mice. Dots per mouse (n=3mice, Welch’s t-test, p=0.1742) **(G)** Percent of all cells in layers 4/5 containing foci in *Cx3cr1^CRE^:Ifnar1^flox/flox^* mice at P5 (n=5mice, Welch’s t-test, p=0.0466) **(H)** Percent of all cells in layers 4/5 containing foci in *Syn1^CRE^:Ifnar1^flox/flox^* mice at P5 (n=3mice, Welch’s t-test, p=0.8939)

### IFN-I is sufficient to promote neuronal engulfment by microglia

To test whether IFN-I was sufficient to drive neuron-engulfing microglia we injected interferon-β (IFN-β) intracerebroventricularly (i.c.v) at P4 and collected tissues at P5 (**Fig. 6A**). We observed robust IFITM3 expression in microglia (86% of microglia) within 300μm of the IFN-β injection, compared to PBS controls (5%; **Fig. 6B-C**). IFITM3+ microglia in IFN-β injected mice were more likely to be phagocytic (**Fig. 6D-E**) and contained almost three times as many phagocytic cups as IFITM3-microglia (**Fig. 6F, Fig. S6B**), mirroring the IRMs we observed in the whisker-deprived context (**Fig. 3D-E**). We also observed about 16% of phagocytic IFITM3+ microglia containing ‘projecting phagosomes’ wrapped around DAPI+ pyknotic nuclei (**Fig. 6E**). In contrast, IFN-β injected mice did not have distended phagosomes or bubble microglia (**Fig. 6G**). We also found that IFN-β injected mice had a significant reduction in neurons with dsDNA breaks (**Fig. 6H-I**). These data suggest that IFN-I promotes accelerated microglial phagocytosis and restricts the accumulation of neurons with dsDNA breaks.

**Figure 6:**
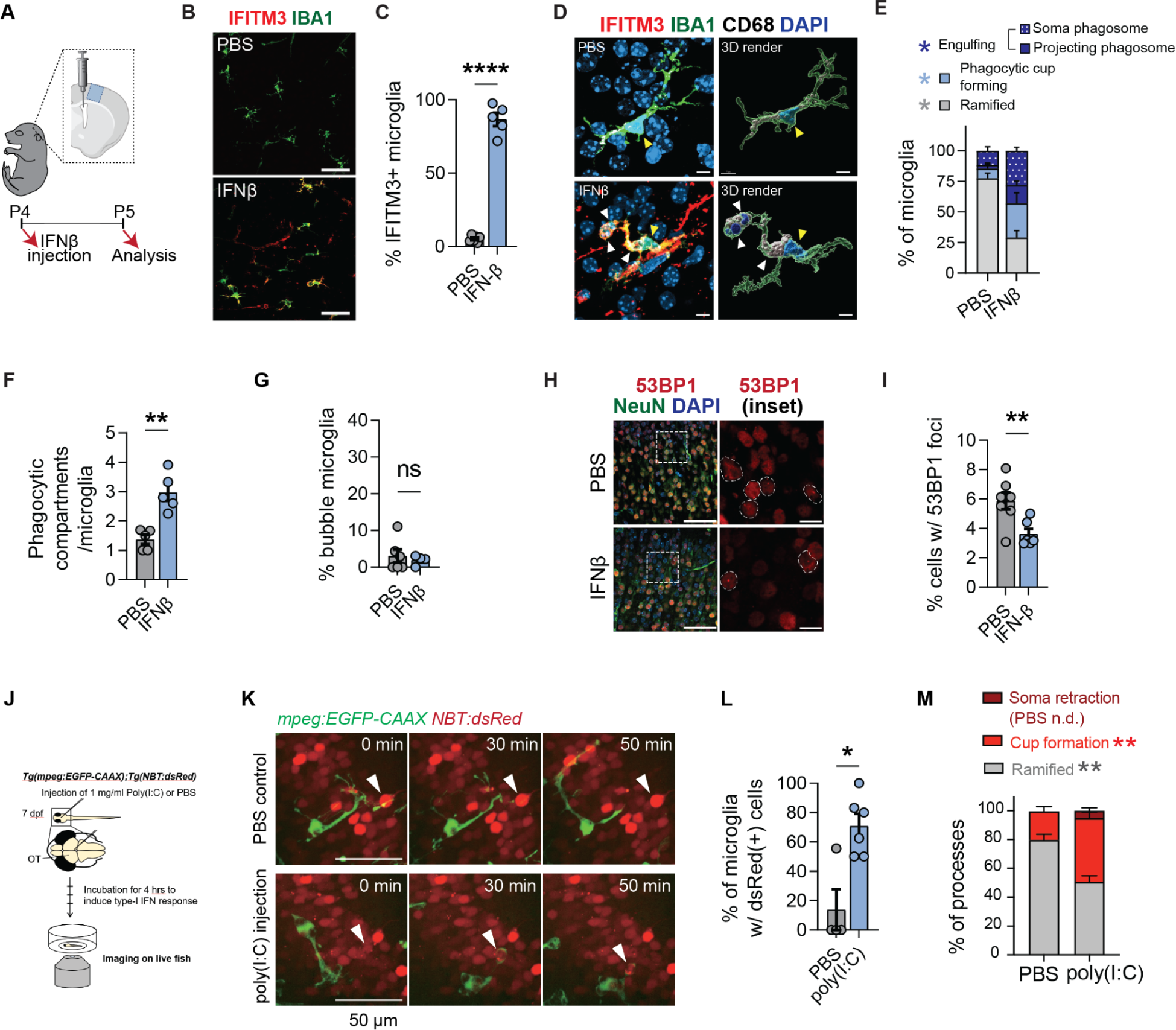
IFN-I is sufficient to promote neuronal engulfment by microglia. **(A)** Diagram of experimental design. 10ng of IFNβ injected at P4. Analysis is performed at P5. Blue square: region analyzed. **(B)** Representative images of PBS vs. IFNβ injected mice motor cortex at the injection site. (scale bar = 50µm) **(C)** IFITM3+ microglia in cortex in PBS vs. IFNβ injected mice. Dots per mouse. Welch’s t-test, p<0.0001, n= 5-6 per group **(D)** Representative images of microglia in PBS vs. IFNβ injected mice with 3D rendering. IFIMT3+ microglia shows both a ‘projecting phagosome’ and a ‘soma phagosome’ (yellow arrow = microglia nucleus, white arrow = DAPI+ containing phagocytic cup, scale bar = 5µm) **(E)** Quantification of microglial morphology subtypes (ramified, w/ phagocytic cup or engulfing with either projecting or soma phagosome) from PBS vs. IFNβ injected mice. Dots per mouse. (n=5-6 per group, 2-way RM ANOVA with Sidak post-hoc test) **(F)** Phagocytic compartments per microglia (IFITM3-vs. IFITM3+) in PBS vs. IFNβ injected mice. Dots per mouse. Welch’s t-test, p=0.0027, n=5 per group. **(G)** Bubble morphology microglia in PBS vs. IFNβ injected mice. Dots per mouse (Welch’s t-test, p=0.5759, n=4-6 mice per group) **(H)** Representative images of 53BP1+ foci containing neurons in PBS vs. IFNβ injected mice motor cortex at the injection site. White dashed line shows inset. Yellow dashed lines outline nuclei with 53BP1+ foci (scale bar = 50µm in low magnification image and scale bar = 10µm in inset) **(I)** Percent of all cells containing 53BP1+ foci in PBS vs. IFNβ injected mice motor cortex at the injection site. Welch’s t-test, p=0.0088, n=6-7 per group. **(J)** Diagram showing parameters for zebrafish poly(I:C) injection and live imaging experiment **(K)** Individual frames (0-50min) from live imaging of green microglia (Tg(mpeg:EGFP-CAAX)) and red neurons (Tg(NBT:dsRed)) in the zebrafish optic tectum after intraventricular injection of PBS or poly(I:C). PBS: the white arrowhead shows a single neuron that is contacted by a microglial process but not engulfed. poly(I:C): the white arrowhead shows a neuron that is contacted, engulfed, and trafficked towards the microglial soma (scale bar = 10μm) **(L)** Quantification showing the percent of microglia engulfing at least one dsRed+ cell during the hour-long video. Dot per fish. (n= 4 PBS, 6 poly(I:C), Welch’s t-test, p=0.161) **(M)** Percent of microglial processes with each of three morphologies (n = 4 PBS, 6 poly(I:C), 2-way ANOVA with Sidak post-hoc test, p=0.1504/0.0056/0.0029)

To temporally define the impact of IFN-I signaling on microglia, we performed live imaging in zebrafish (*Danio rerio*; **Fig. 6J**). This is an ideal model system for imaging the intact developing brain, and contains an evolutionarily conserved antiviral IFN-I system^59,60^. A dedicated subset of microglia in the zebrafish optic tectum engulfed neuronal soma during development^58,61^. We activated IFN-I signaling 4 hours prior to imaging with an intraventricular injection of the viral mimic poly(I:C) into zebrafish expressing the myeloid reporter *Tg(mpeg:EGFP-CAAX)* and the pan-neuronal reporter *Tg(NBT:dsRed*) (**Fig. 6J**), which includes a resident population of neuron-engulfing microglia^61,62^. IFN-I stimulation significantly increased phagocytic cup formation and accelerated the enveloping, engulfment, and retraction of neuronal soma (**Fig. 6K, Fig. 6M, Fig. S6D**, **Supplemental Movies 2-3**). Poly(I:C) led to a four-fold increase in the percent of microglia engulfing at least one neuron (**Fig. 6L**) and increased the percentage of microglia engulfing multiple neurons (**Fig. S6C**), similar to what we observed in IFN-responsive murine microglia after whisker deprivation (**Fig. 3F-H**). Thus IFN-I signaling is sufficient to induce microglial engulfment of neurons in both murine and zebrafish models.

### IFN-I responsive microglia restrict tactile hypersensitivity

We next examined the impact of IFN-I signaling on somatosensory cortical development and function. As we had observed that IRMs preferentially engulf L5 neurons during early life, we quantified excitatory and inhibitory neuronal subsets in *Ifnar1^-/-^* mice vs. littermate controls at P15, once neuronal numbers are stabilized^63^. We found a significant increase in the density of CTIP2+ excitatory neurons, which are enriched in L5/6 (**Fig. 7A, E**), whereas the pan-cortical excitatory marker SATB2 was not altered (**Fig. 7B, F**). Conversely, we found a significant decrease in the density of PV+ inhibitory neurons, which are enriched in L4/5 (**Fig. 7C, G**), and a trend towards fewer SST+ interneurons (**Fig. 7D, H**). These data suggest that IFN-I deficiency leads to increased ratio of excitatory vs. inhibitory neurons in deeper cortical layers. This is consistent with defective elimination of some excitatory neurons, although a more general impact on cortical maturation cannot be ruled out, as markers like CTIP2 and PV can be maturation-dependent in some contexts^64^.

**Figure 7:**
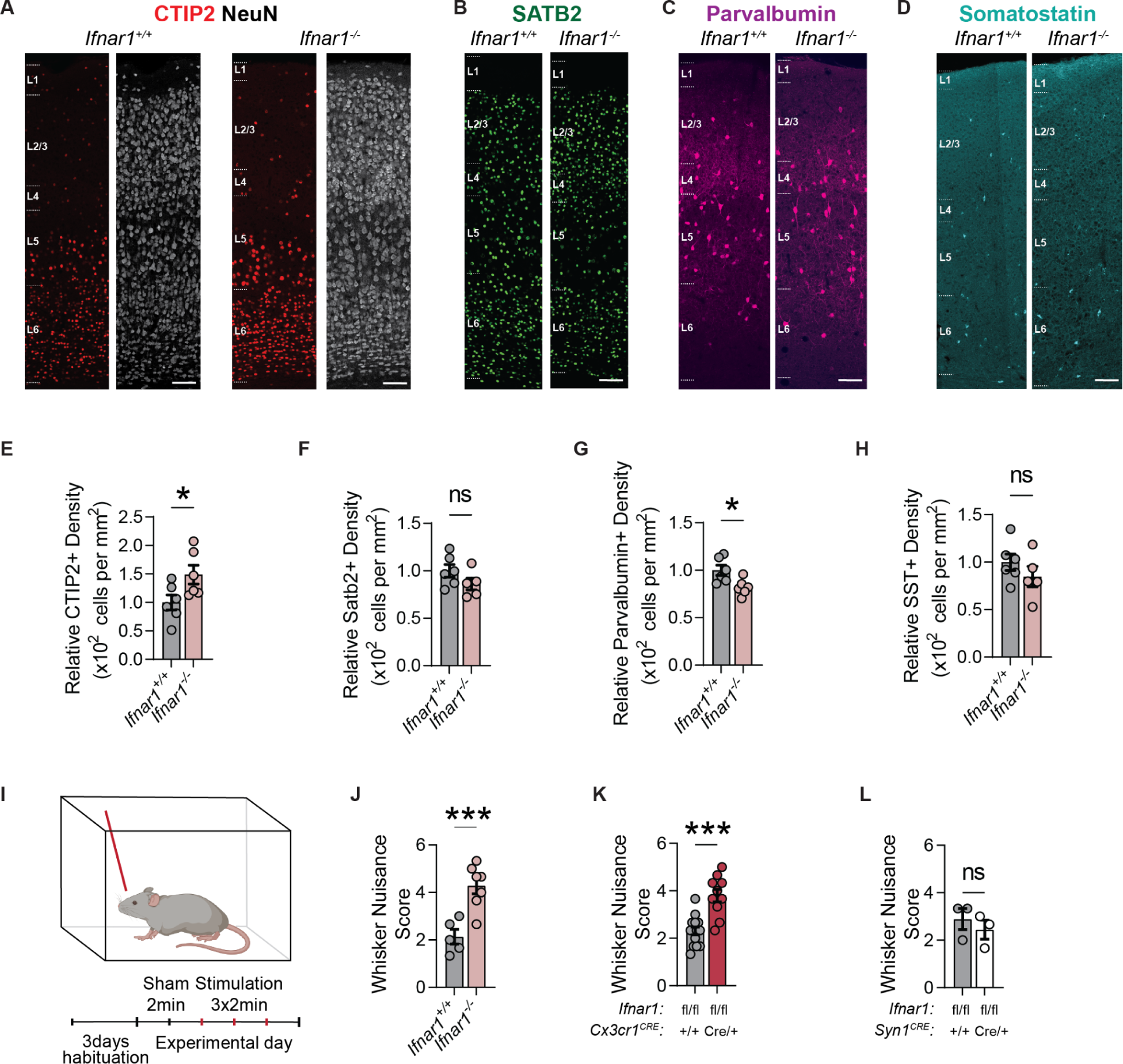
IFN-I responsive microglia prevent tactile hypersensitivity. **(A)** Representative images of CTIP2 in the somatosensory cortex in P15 *Ifnar1^+/+^* and *Ifnar1^-/-^* mice. White dashed lines highlight the cortical layers (scale bar = 100μm) **(B)** Representative images of SATB2 in the somatosensory cortex in P15 *Ifnar1^+/+^* and *Ifnar1^-/-^* mice. (scale bar = 100μm) **(C)** Representative images of Parvalbumin in the somatosensory cortex in P15 *Ifnar1^+/+^* and *Ifnar1^-/-^* mice. (scale bar = 100μm) **(D)** Representative images of Somatostatin in the somatosensory cortex in P15 *Ifnar1^+/+^* and *Ifnar1^-/-^* mice. (scale bar = 100μm) **(E)** Relative CTIP2+ neuron density across cortical layers at P15 *Ifnar1^+/+^* and *Ifnar1^-/-^* mice. Data are represented as mean±sem. Dots=mice. Welch’s t-test, p=0.0443, n=6 mice per group. **(F)** SATB2+ neuronal density across cortical layers in P15 *Ifnar1^+/+^* and *Ifnar1^-/-^* mice. Data are mean±sem. Dots = mice. Welch’s t-test, p=0.1638, n=5-6 mice per group. **(G)** Parvalbumin+ neuronal density across cortical layers in P15 *Ifnar1^+/+^* and *Ifnar1^-/-^* mice. Data are mean±sem. Dots = mice. Welch’s t-test, p=0.0197, n=6 mice per group. **(H)** Somatostatin+ neuronal density across cortical layers in P15 *Ifnar1^+/+^* and *Ifnar1^-/-^* mice. Data are mean±sem. Dots = mice. Welch’s t-test, p=0.3038, n=5-6 mice per group. **(I)** Schematic of whisker nuisance assay. **(J)** Whisker nuisance score in P15 *Ifnar1^+/+^* and *Ifnar1^-/-^* mice. Data are mean±sem. Dots = mice. Welch’s t-test, p=0.0009, n=5-7 mice per group, 3 independent experiments. **(K)** Whisker nuisance score in P15 littermate control (*Ifnar1^flox/flox^*) vs. *Cx3cr1^CRE^:Ifnar1^flox/flox^* mice. Data are mean±sem. Dots = mice. Welch’s t-test, p=0.0002, n=10-14 mice per group, 4 independent experiments. **(L)** Whisker nuisance score in P15 littermate control (*Ifnar1^flox/flox^*) vs. *Syn1^CRE^:Ifnar1^flox/flox^* mice. Data are mean±sem. Dots = mice. Welch’s t-test, p=0.4993, n=3 mice per group.

We next examined the impact of IRMs on somatosensory function. We used a whisker nuisance assay to quantify behavioral responses to tactile stimulation in juvenile mice (P15)^65–67^. Following habituation to context and probe, mice underwent three brief episodes of unilateral whisker stimulation (**Fig. 7I**, see Methods). Tactile sensitivity to whisker stimulation was tabulated via manual visual scoring of single behaviors (see methods^66^). Both global and microglial-specific deletion of *Ifnar1^-/-^* led to a significant increase in tactile hypersensitivity as reflected by aggression and increased avoidance towards the probe (**Fig. 7J-K**). In contrast, conditional deletion of *Ifnar1* in neurons had no impact on tactile responses (**Fig. 7L**). These data directly demonstrate a role for IRMs in restricting tactile hypersensitivity.

## Discussion

IFN-I responses are central to antiviral defense but their role in physiology is comparatively unknown^68–70^. Here we describe a Type I interferon responsive microglial subset that is required for healthy brain development. These cells were rare but detectable during typical development, but deficient microglial IFN-I responses resulted in a large fraction of microglia with phagocytic dysfunction, accumulation of damaged neurons, and alterations in cortical layer development and behavior. These data suggest that IRMs may be a transient but critical microglial state during brain development. IRMs expanded 20-fold after whisker deprivation, and were frequently digesting one cell while in the process of enclosing another. It is possible that these microglia could boost phagocytosis in settings where neuronal turnover is high. While a similar IRM signature is conserved across multiple viral as well as sterile contexts, whether they have similar functional roles in these settings remains to be determined.

The finding that accumulation of dsDNA-damaged neurons is inversely correlated with the presence of IFN-I responsive microglia strongly suggests that intact IFN-I signaling in the developing brain optimizes the efficient elimination of damaged neurons. Determining the cellular sources of IFN-α/β and the neuronal cues that engage a microglial IFN-I response are essential future questions. One potential model is that exposure to damage associated molecular patterns (DAMPs) from stressed cells triggers conserved responses adapted to detect nucleic acids^71–74^, triggering an autocrine loop in microglia which enhances phagocytic efficiency via IFN-α/β release. In this model, the tightly regulated physiological engulfment of neurons could become pathological if high IFN-I tone or abundant nucleic acids trigger a more widespread microglial response and lead to indiscriminate cell death and neuroinflammation.

Epidemiologic links between maternal infections to neurodevelopmental disorders such as schizophrenia and autism spectrum disorders (ASD)^75^ have inspired the “maternal immune activation” (MIA) model of neurodevelopmental disorders, which usually relies on maternal delivery of the potent IFN-I agonist poly I:C^76^. Deficits in sensory processing are common in ASD^77^ and autism risk genes expressed exclusively in somatosensory neurons can lead to ASD-like behavioral deficits, suggesting a key role for somatosensory processing in the overall ^78,79^. Our data suggest that loss of IFN-I signaling results in prominent tactile hypersensitivity, which is also observed in animal models of ASD-like behaviors^80^. Further defining whether viral infections might drive the emergence of IFN-I responsive microglia and promote neuronal engulfment could reveal mechanistic links between developmental immune activation and neuropsychiatric diseases.

## Supporting information

Supplemental_Movie_1-Bubble

Supplemental_Movie_2_PBS_MG_video

Supplemental_Movie_3_PolyIC_MG_video

Supplementary Table 2- MACS_allcells_clustermarkers

Supplementary Table 3 - Microglia_only_allclustermarkers

Supplementary Table 4 -bulk_models_Legend

Supplementary Table 5 - singlecellmodels

Supplementary Table 6 - Spliced_unsplicedratio

---

**FIGURE S1:**
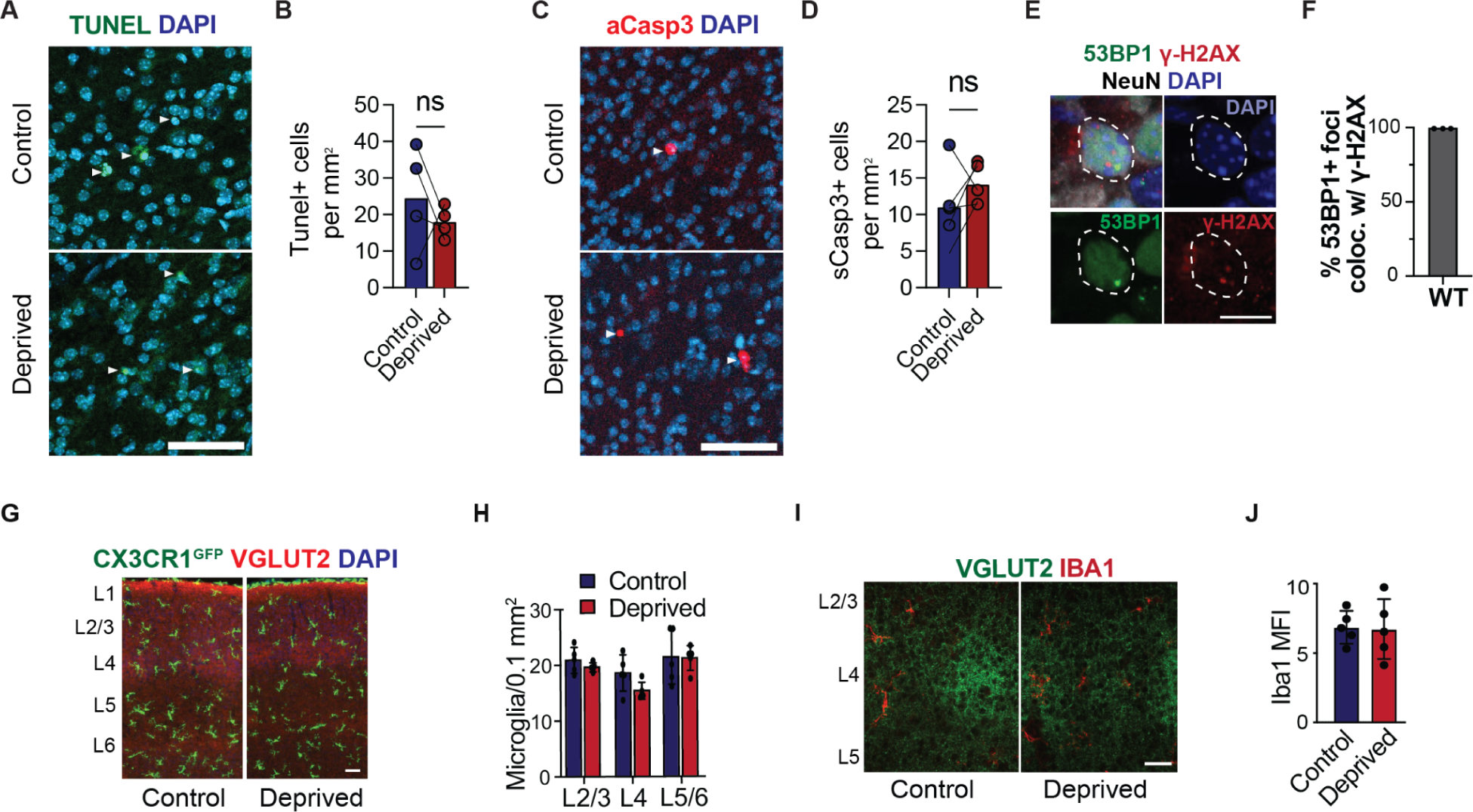
Additional characterization of the partial whisker deprivation model, supplemental to Figure 1. **(A)** Representative image of Tunel+ cells in barrel cortex of control vs. deprived hemispheres (scale bar = 50μm) **(B)** Density of Tunel+ cells in the barrel cortex of control vs. deprived hemispheres per mm^2^. Dots=mice, lines connect both hemispheres from the same mouse. (paired t-test, n=4 mice, p=0.4999) **(C)** Representative image of aCaspase3+ cells in barrel cortex of control vs. deprived hemispheres (scale bar = 50μm) **(D)** Density of aCaspase3+ cells in the barrel cortex of control vs. deprived hemispheres per mm^2^. Dots=mice, line connect both hemispheres from the same mouse. (paired t-test, n=5 mice, p=0.3087) **(E)** Representative images showing colocalization of ɣH2AX and 53BP1 foci within a neuronal nucleus. (scale bar = 5μm) **(F)** Quantification of percent 53BP1 foci+ cells that also had ɣH2AX+ foci. (n=3 mice) **(G)** Representative image of microglia (Cx3cr1:GFP+) in P7 cortex (scale bar = 20μm) **(H)** Microglial density across cortical layers (n =4 mice) **(I)** Representative images of IBA1 intensity through the barrel cortex at P7. (scale bar = 25μm) **(J)** Quantification of IBA1 intensity through the barrel cortex at P7 (n = 5 mice)

**FIGURE S2:**
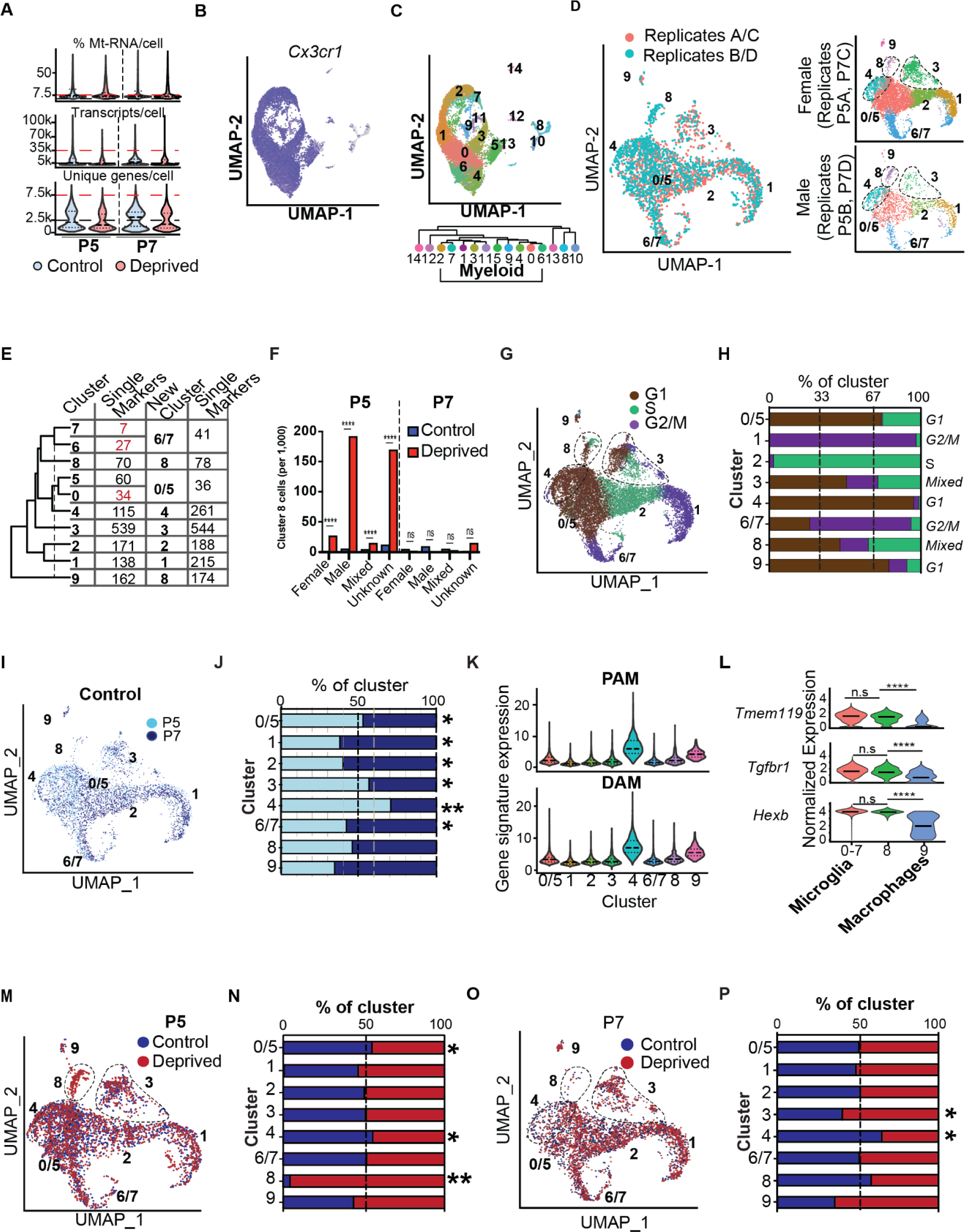
Quality control and additional analyses of single cell data, supplemental to Figure 1. **(A)** Quality control metrics for microglial single cell sequencing. “% Mt-RNA/cell” = mitochondrial RNA content per cell. Dashed lines indicate minimum and maximum threshold settings. Dotted lines inside violin plots represent median, 1^st^ and 3^rd^ quartiles **(B)** Feature plot of *Cx3cr1* expression. **(C)** Unsupervised clustering of all thresholded cells. Clustering tree show selected *Cx3cr1+* cells used in all subsequent analyses. **(D)** Comparison of biological replicates bioinformatically segregated via expression of male-specific (*Ddx3y* and *Eif2s3y*) and female-specific (*Xist* and *Tsix*) transcripts. Female and Male replicates also shown colored by cluster in UMAP space. (Replicate A: 3 female mice at P5; Replicate B: 3 male mice at P5; Replicate C: 3 female mice at P7; Replicate D: 1 male mouse at P7) **(E)** Table showing the number of uniquely upregulated genes (lfc>0.15,p.adj<10^-5^) before and after combining clusters 6/7 and 0/5.The dotted line shows clustering threshold used. The resultant clusters were not closely related to any other clusters with few unique DEGs. **(F)** Cluster 8 enrichment after whisker deprivation in definitively female (*Xist*/*Tsix*+; *Ddx3y*/*Eif2s3y*-), definitively male (*Ddx3y*/*Eif2s3y*+; *Xist*/*Tsix*-), mixed (*Xist*/*Tsix*+; *Ddx3y*/*Eif2s3y*-) and unknown (*Xist*/*Tsix*-; *Ddx3y*/*Eif2s3y*-) cells. (Chi-square test with Bonferroni correction, ****pAdj<10^-10^) **(G)** UMAP plot showing cell cycle phase assignment determined using annotation from *Kowalczyk et al* ^82^. **(H)** Quantification of cluster composition by cell cycle phase. X-axis=%cells per cluster in G1, S, or G2/M. Labels to the right indicate that cluster’s predominant phase. **(I)** UMAP plot of P5 and P7 control cells, normalized for abundance **(J)** Quantification of cluster composition by age in control hemispheres. X-axis represents percent of cells in each cluster from P5 (light blue) or P7 (dark blue) mice, normalized for total number of cells per sample. (Chi-square test with Bonferroni correction, *pAdj<0.01, **pAdj<10^-25^) **(K)** DAM gene signature expression by cluster. The signature represents the 83 genes upregulated by DAMs that were also expressed in this dataset (lfc > 1.5, adj. p <10^-8^)^29^. Dotted lines represent median and 1^st^ and 3^rd^ quartiles. PAM gene signature expression by cluster. The signature represents the 42 genes conserved between two published datasets of a developmental PAM signature that were also expressed in this dataset (lfc > 1.5, adj. p <10^-8^)^27,28^. Dotted lines represent median and 1^st^ and 3^rd^ quartiles **(L)** Canonical microglial gene expression across microglial clusters 0-6 (pooled), microglial cluster 8, and putative macrophage cluster 9. Line =median. (MAST DE test, ****p<10^-25^) **(M)** UMAP of P5 cells colored by condition (control vs. whisker deprived) **(N)** Quantification of P5 cluster composition by condition. X-axis represents % cells in each cluster from the control (blue) or deprived (red) hemispheres, normalized for total number of cells per sample. (Chi-square test with Bonferroni correction, *pAdj<0.01, **pAdj<10^-25^) **(O)** UMAP of P7 cells colored by condition (control vs. whisker deprived) **(P)** Quantification of P7 cluster composition by condition. X-axis represents percent of cells in each cluster from the control (blue) or deprived (red) hemispheres, normalized for total number of cells per sample. (Chi-square test with Bonferroni correction, *pAdj<0.01, **pAdj<10^-25^)

**Figure S3:**
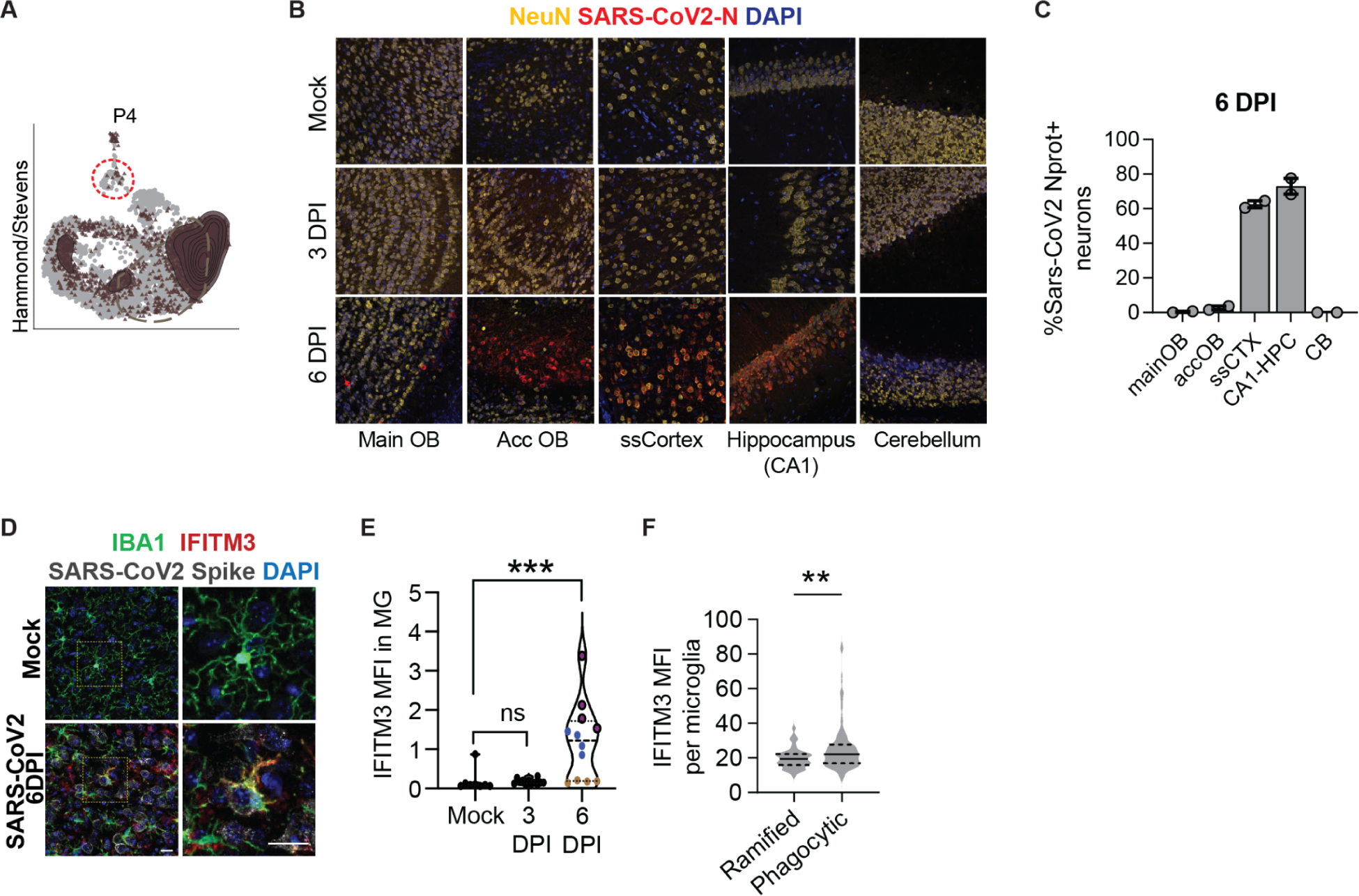
Additional characterization of Sars-CoV2 infection model, supplemental to Figure 2. **(A)** The microglial P5/P7 dataset (colored dots) was re-clustered to create a reference PCA and UMAP map to which other datasets (grey dots) were aligned. Brown triangles represent the external dataset from whole brain P4 in Hammond et al.^27^, red dotted circle highlight cluster 8, grey dashed circle highlight homeostatic cluster 0/5. **(B)** Representative images of 5 brain regions after mock infection, and 3 or 6 DPI, labeled for SARS-CoV-2 N-protein and NeuN (neurons). Main OB: Olfactory Bulb, Acc OB: Accessory Olfactory Bulb, Cortex: somatosensory cortex. **(C)** Number of SARS-CoV-2 N-protein+ cells in the indicated brain regions 6DPI mice. Dots per mouse (n= 2 mice) **(D)** Representative images in somatosensory cortex showing IFITM3 staining in IBA1+ microglia from mock vs. SARS-CoV-2 infected mice at 6 DPI (right = inset, scale bar =20μm) **(E)** Quantification of IFITM3 mean fluorescence intensity within microglia. Each color represents an individual mouse (n=2 mock-infected mice, 3 each at 3 and 6 DPI, Kruskal-Wallis test, p =0.1961/0.0001) **(F)** Quantification of IFITM3 intensity in phagocytic and non-phagocytic (ramified) cells from 6DPI mice. 25^th^, 50^th^, and 75^th^ percentiles shown as dotted lines. Dots represent cells (Welch’s t-test, p=0.0003, n= 40-145 cells from 2 mice)

**Figure S4:**
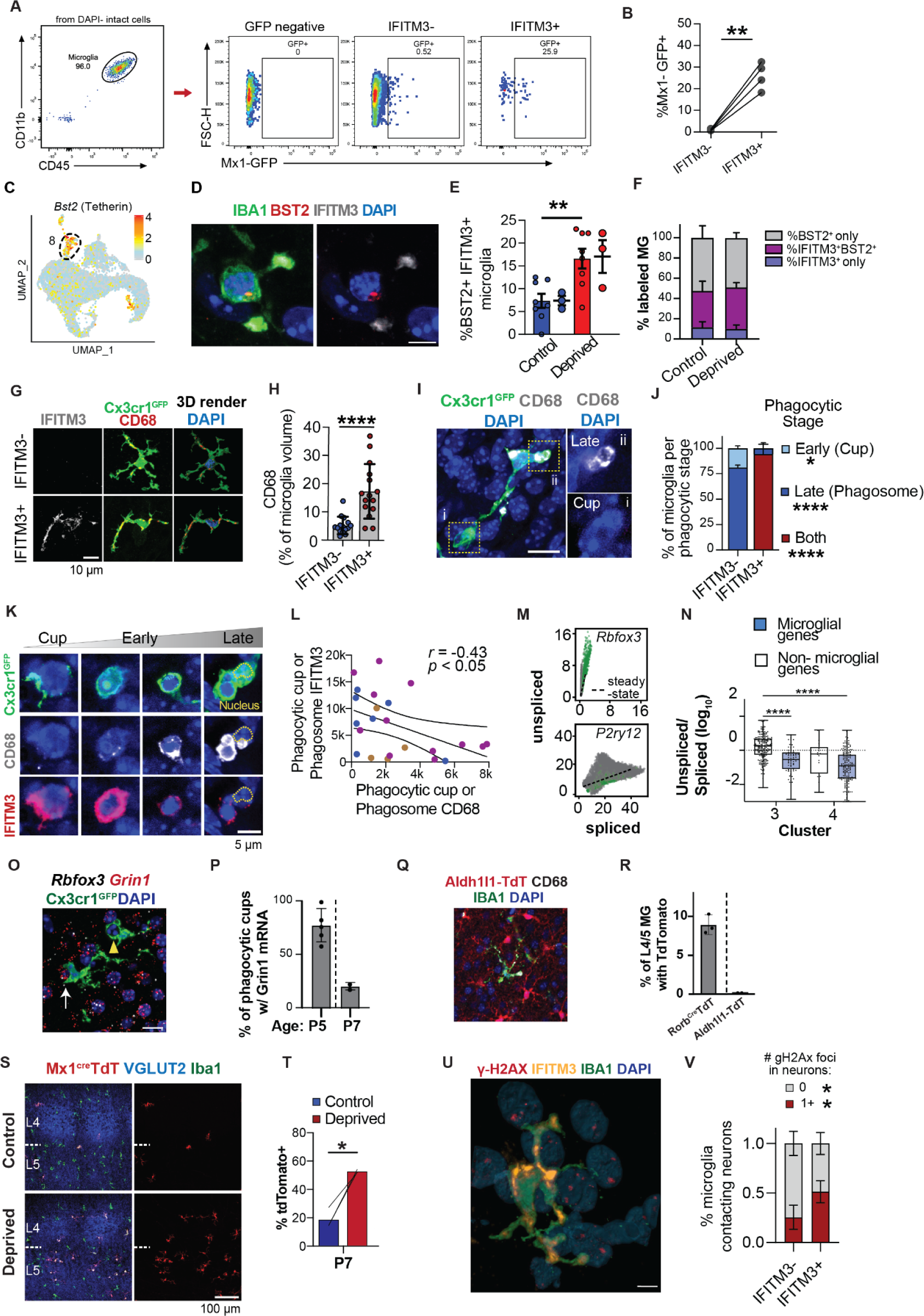
Additional characterization of neuron-engulfing microglia during cortical development, supplemental to Figure 3. **(A)** Representative gating of IFITM3 expression in CD11b^+^/CD45^low^ microglia in control and deprived hemispheres (FMO: fluorescence minus one, negative control) and representative flow gating of *Mx1^GFP+^* microglia from IFITM3-vs. IFITM3+ populations **(B)** Mx1^GFP+^ microglia within IFITM3+ and IFITM3-populations. Lines connect values from the same mouse (n = 4 mice, paired t-test. p=0.0025) **(C)** Feature plot showing normalized expression of *Bst2* in Cluster 8 microglia **(D)** Colocalization of BST2/Tetherin, IFITM3, and microglia in deprived barrel cortex (scale bar = 10μm) **(E)** Percent of microglia co-expressing BST2 and IFITM3 (n = 8 images per condition from 3 mice, Welch’s t-test, p=0.0043) **(F)** Percent of IBA1+ microglia expressing either IFITM3 only (blue), BST2/Tetherin only (grey) or both (purple), excluding double negative microglia (n=3 mice) **(G)** Representative images and 3D-rendering of CD68+ lysosomes within IFITM3+ and IFITM3-microglia in L4 (scale bar = 10μm) **(H)** Quantification of % CD68 volume of total microglial volume in IFITM3+ and IFITM3-microglia (15 IFITM3-and 15 IFITM3+ from n= 3mice, Mann-Whitney test, p<0.0001) **(I)** Representative image of a Cx3cr1^GFP+^ microglia highlighting a phagocytic cup (i; CD68-, non-pyknotic DAPI signal, incomplete DAPI envelopment by microglial processes) and a late phagosome or phagolysosome (ii; CD68+, pyknotic DAPI signal, complete envelopment by microglial processes) (scale bar = 10μm) **(J)** Distribution of early phagocytic cups and late phagosomes as a proportion, in microglia containing at least one phagosome (n =7-10 cells each from 3 mice/group, 2-way RM ANOVA with Sidak post-hoc test, p=0.0313/<0.0001/<0.0001) **(K)** Representative images showing changes in IFITM3 and CD68 expression at different stages of phagocytosis (yellow dotted line = microglial nucleus, scale bar = 5μm) **(L)** Correlation of CD68 and IFITM3 mean fluorescence intensity per phagocytic compartment, (n = 23 cells from 3 mice, r Spearman correlation coefficient = −0.43, p<0.05) **(M)** Representative plots of transcript splicing state for a canonical neuronal gene (*Rbfox3*, which encodes NeuN) vs. a microglial gene (*P2ry12*). Each dot represents one cell; units are in transcript counts per cell. Green dots=Cluster 3; grey dots are cells in all other clusters. **(N)** Ratio of unspliced to spliced transcripts (log_10_) for genes upregulated in Clusters 3 and 4, for canonical microglial genes (blue) vs. genes enriched in other cell types (white). “Enriched” =10x higher FPKM in microglia than the mean of other cell types from^83^. Plots show range, median, and first to third quartiles. (Welch’s ANOVA with Tamhane’s T2 test for multiple comparisons, p<0.0001). **(O)** Representative image of Cx3cr1^GFP+^ microglia engulfing a DAPI+ nucleus expressing neuronal mRNA transcripts *Rbfox3* and *Grin1*. White arrow: a microglial phagocytic cup. Yellow arrowhead: enclosed phagosome lacking neuronal transcripts. (Scale bar = 10μm). **(P)** Percent of microglial phagocytic cups enveloping nuclei containing *Grin1* transcripts at the specified ages. Dots represent mice (n= 5 mice at P5 and 2 mice at P7) **(Q)** Representative image of IBA1+ microglia and Aldh11L1-TdT+ astrocytes in barrel cortex. (scale bar = 50μm) **(R)** Quantification of percent microglia containing TdTomato signal in Rorb^cre^:TdT or Aldh1l1^TdT^ transgenic mice. Dots=mice. **(S)** Representative images of tdTomato+ cells (*Mx1-Cre;Rosa26-LSL-TdT*) in L4/5 of the barrel cortex co-labeled with microglial marker IBA1 and VGLUT2 to highlight barrels. (Scale bar= 100μm) **(T)** TdTomato+ microglia in control and deprived cortices. Lines connect control and deprived hemispheres from the same mouse (n = 3 mice, paired t-test, p=0.0186) **(U)** 3D rendering of IFITM3+ IBA1+ microglia contacting cells with ɣH2Ax+ foci in WT deprived cortex. **(V)** Mean number of ɣH2Ax+ foci in cells contacted by IFITM3-vs. IFITM3+ microglia in WT deprived cortex. Dots per mouse. Lines connect values from the same mouse (Paired t-test, p=0.0290, n=17 vs 23 microglia from n=5 mice)

**Figure S5:**
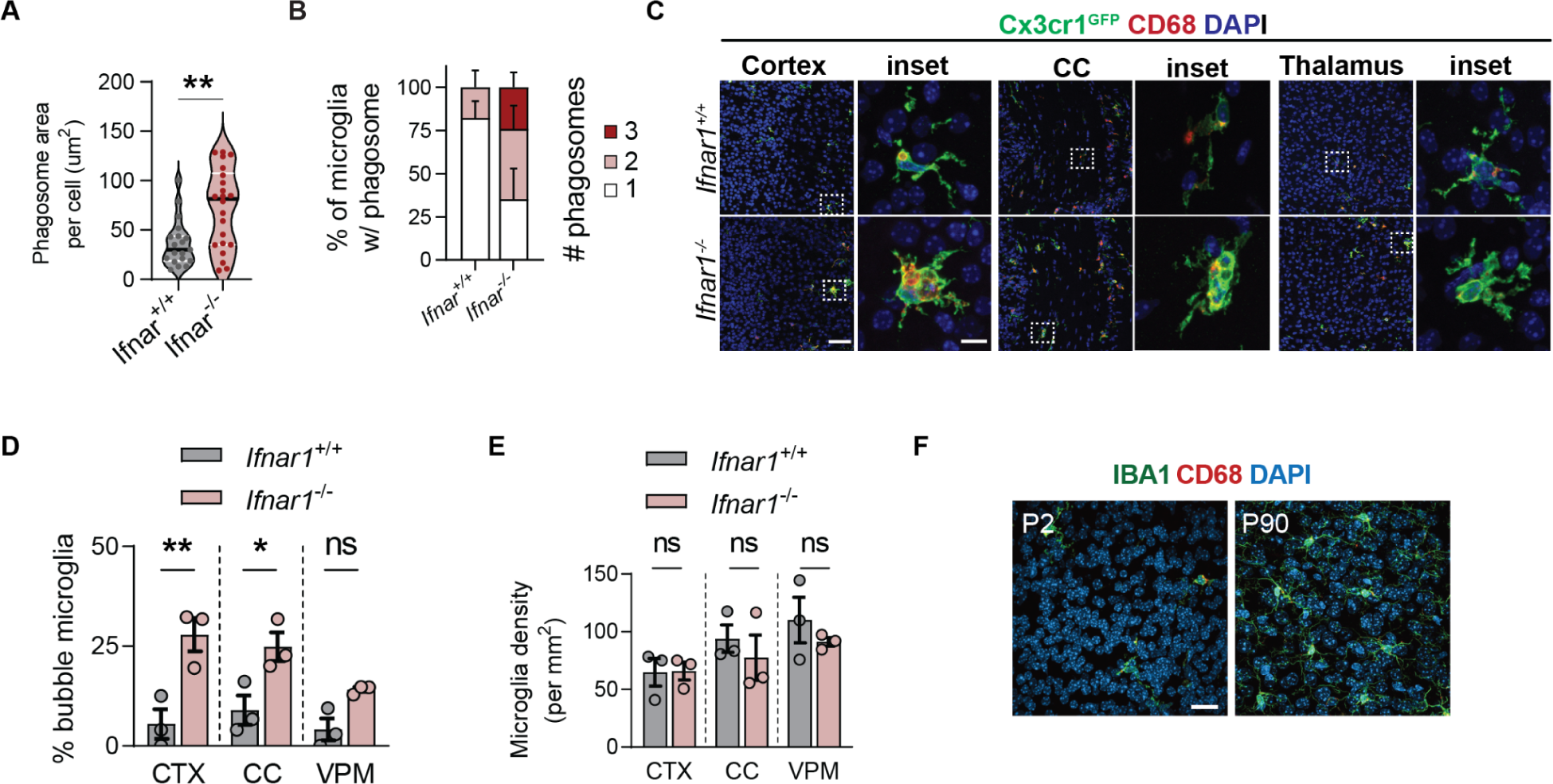
Additional characterization of *Ifnar1^-/-^* mice across brain regions and developmental stages, supplemental to Figure 4. **(A)** Total area of phagocytic compartments per microglia in L4/5 barrel cortex from *Ifnar1^+/+^* and *Ifnar1^-/-^* mice. Dots per microglia. Mann-Whitney test, p=0.0053, n=19-22 microglia from 3 mice per group **(B)** Number of phagocytic compartment per microglia in *Ifnar1^+/+^* and *Ifnar1^-/-^* mice at P5 (n=3 mice) **(C)** Representative images showing microglial morphology in different brain regions (scale bar = 50 μm (left, low power), 10 μm (inset, high power)) **(D)** Percent bubble microglia as a percent of total microglia per brain region in *Ifnar1^+/+^* and *Ifnar1^-/-^* mice (n=3 mice per genotype, 2-way ANOVA with Sidak’s post-hoc test, p=0.0013/0.0154/0.1507) **(E)** Microglial density in in *Ifnar1^+/+^* and *Ifnar1^-/-^* mice by brain region (n=3 mice per genotype, 2-way ANOVA with Sidak’s post-hoc test, p= >0.9999/0.8004/0.7291) **(F)** Representative confocal images of IBA1+ microglia in L4/5 barrel cortex WT mice at P2 and P90 quantified in Fig. 4H

**Figure S6:**
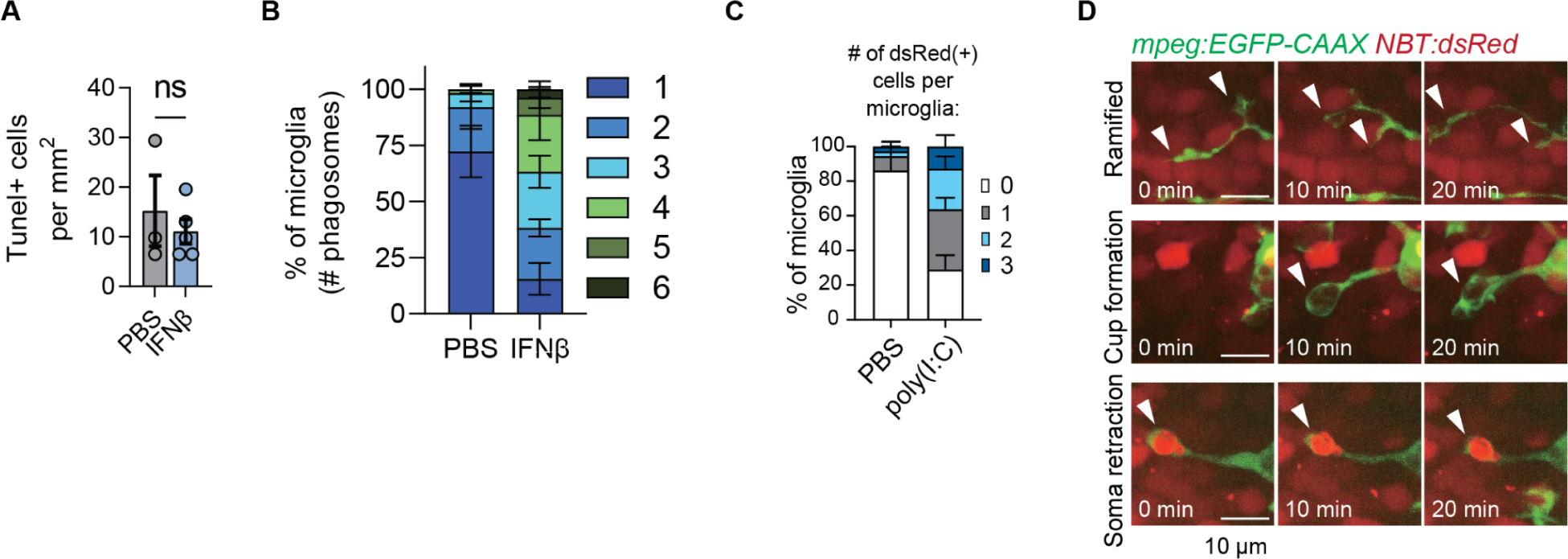
Additional characterization of the gain of function models in mouse and zebrafish, supplemental to Figure 6. **(A)** Density of Tunel+ cells per mm^2^ in PBS vs. IFNβ injected mice motor cortex at the injection site. Dots per mouse. Welch’s t-test, p=0.6289, n=3-5 per group. **(B)** Percent of microglia containing 1 to 6 phagosomes per cell, among microglia with at least one phagosome (n = 5 mice per group, 36-37 cells per group) in PBS vs. IFNβ injected mice motor cortex at the injection site **(C)** Representative images defining the following microglial morphologies: ramified, cup formation and soma retraction. White arrowheads show interactions between the microglial process (mpeg:EGFP-CAAX) and surrounding neuronal cell bodies (NBT:dsRed) (scale bar = 10μm) **(D)** Percent of microglia engulfing 0, 1, 2, or 3 dsRed+ cells during image acquisition. (n = 4 PBS, 6 poly(I:C))

**Supplemental movie 1:** Imaris reconstructed video scanning through the z-axis of a dysmorphic bubble microglia shown in Fig. 4F. Green = IBA1 (microglia), blue = DAPI (nucleus), red = CD68 (lysosomes).

**Supplemental movies 2-3:** Representative live imaging videos of microglia (*Tg(mpeg:EGFP-CAAX))* and neurons (*Tg(NBT:dsRed)*), from vehicle injected (Supplemental movie 2) or polyI:C injected (supplemental movie 3) Each frame consists of one 50 mm z-stack acquired once every 5 minutes, and each video was acquired over one hour.

**Table S1:** Sample sizes and cell yields from the single cell sequencing in **Fig.1**

| Sample age | Sample description | #cells after QC |
| --- | --- | --- |
| **Microglial only RNA seq (isolated by dounce homogenization), Fig. 1** |  |  |
| P5 | 3 Females, pooled, control and deprived hemispheres | 1360 control, 1265 deprived |
| P5 | 3 Male, pooled, control and deprived hemispheres | 1595 control, 1290 deprived |
| P7 | 3, Females pooled, control and deprived hemispheres | 2929 control, 2224 deprived |
| P7 | 1 Male, control and deprived hemispheres | 1165 control, 502 deprived |

**Table S2 (excel file):** Differentially expressed genes for CD11b+ single-cell sequencing clustering shown in Fig. 3J.

**Table S3 (excel file):** Differential expression analysis for in-silico sorted macrophages isolated at P5 and P7, see also Fig. 1H, Fig. K-L, Fig. 3J.

**Table S4 (excel file):** Extended legend for **Fig. 2A** including references for all publicly available bulk microglial sequencing datasets used. Table adapted from Friedman et al 2018^30^, figure 4.

**Table S5 (excel file):** Extended legend for **Fig. 2B-C**, including references for all publicly available published single cell sequencing datasets used.

**Table S6 (excel file):** Ratio of spliced to unspliced transcripts identified per gene calculated using velocyto and scvelo on the P5/P7 microglial single cell RNAseq dataset. See **Fig. 3I**.

**Table S7 (excel file):** Summary of statistical analyses.

## Acknowledgements

We are grateful to members of the Molofsky Lab for helpful comments on the manuscript, Dr. Rafael Han for assistance with tissue preparation, and Dr. Ari Molofsky for helpful feedback on the manuscript. Thanks to the Chan-Zuckerberg Biohub for sequencing support. Funding: A.V.M is supported by the Pew Charitable Trusts, NIMH (R01MH119349 and DP2MH116507), and the Burroughs Welcome Fund. T.J.N. was supported by NIMH RF1MH121268. L.C.D. received support from the Matilda Edlund Scholarship and the Genentech Fellowship. P.T.N. was supported by a graduate research fellowship from the National Science Foundation (Grant #1650113). C.C.E. was supported by a gift from Marilyn Waldman and T32 training grant (5T32AI007334). C.C. was supported by NIA RF1AG061874.

## Author contributions

Conceptualization: A.V.M, L.C.D., C.C.E. and P.T.N.; Methodology, L.C.D., A.V.M, P.T.N., I.D.V., C.C.E., H. N-I., Y.X., R.A. B.S., H.N., N.J.S., T.N.; Investigation: L.C.D., P.T.N., I.D.V, S.E.T., C.C.E, E.W., Y.X., P.V.L., C.L.L., B.C., H.N., N.J.S., S.R.A.; Writing – Original Draft, L.C.D., P.T.N., C.C.E., and A.V.M.; Writing – Review & Editing, all co-authors; Funding Acquisition, A.V.M. Resources, A.V.M., R.A., Supervision, A.V.M., T.N., R.A., B.S., and I.D.V.

### Data availability

Supplement contains additional data. All data needed to evaluate the conclusions in the paper are present in the paper or the Supplementary Materials. Searchable database available at https://www.annamolofskylab.org/microglia-sequencing. RNA sequencing data is available through GEO at <u>GSE173173</u>. Any additional data needed to evaluate the paper will be provided upon request.

### Code availability

R and Python code used to analyze single cell datasets is available on GitHub at https://github.com/lcdorman/IFNresponseCode.

## FULL METHODS

−CONTACT FOR REAGENT AND RESOURCE SHARING

Anna Molofsky

## −EXPERIMENTAL MODELS AND SUBJECT DETAILS

### Mice

All mouse strains were maintained in the University of California San Francisco specific pathogen–free animal facility, and all animal protocols were approved by and in accordance with the guidelines established by the Institutional Animal Care and Use Committee and Laboratory Animal Resource Center. Mice were housed in a 12-hour light/dark cycle (7 am-7pm) at 68-79° F and 30-70% humidity. Littermate controls were used for all experiments when feasible, and reporter mice were backcrossed >10 generations on a C57Bl/6 background (Cx3cr1-GFP) or Swiss-Webster (Aldh1l1-eGFP). The following mouse strains used are described in the table below and are as referenced in the text: Cx3cr1-GFP (JAX #00582); Aldh1l1-eGFP (Gensat); Ifnar1^-/-^ (JAX #028288); Mx1-GFP (JAX #033219); Mx1-Cre (JAX #003556); Ai14 (JAX #007908); B6.Cg-Tg(K18-ACE2) 2Prlmn/J (JAX #034860); Rorb-Ires2-Cre-D (Jax 023526); Aldh1l1-tdTomato (MMRRC, RRID:MMRRC_036700-UCD); B6.Cg-Tg(Syn1-cre)671Jxm/J (JAX #003966); Ifnar1 flox B6(Cg)-Ifnar1tm1.1Ees/J (JAX #028256); Cx3cr1-Cre (MMRRC, RRID:MMRRC_036395-UCD).

### Zebrafish

Fish were maintained in recirculating habitats at 28.5 °C and on a 14/10-h light/dark cycle. Embryos were collected after natural spawns and incubated at 28.5 °C. Larvae were imaged at 7 days post fertilization (dpf), a developmental stage at which sex cannot be determined. In this study, we used the double transgenic fish *Tg(mpeg:EGFP-CAAX);Tg(NBT:dsRed)* (zfin ID: ZDB-TGCONSTRCT-191211-1 and ZDB-TGCONSTRCT-081023-2) on a casper background to visualize microglia and neurons. All zebrafish protocols were approved by and in accordance with the ethical guidelines established by the UCSF Institutional Animal Care and Use Committee and Laboratory Animal Resource Center (LARC).

## −METHOD DETAILS

### Whisker lesions

Whisker ablations were performed under hypothermia-induced anesthesia and followed by topical application of lidocaine for pain management. Postnatal day two pups were anesthetized for 3 minutes in ice. An incision was made in the whisker pad along whisker rows B and D of one side of the face, and silver nitrate was used to cauterize the exposed whisker follicles in that row. After topical 2% lidocaine application the mice were reacclimated on a heating pad for at least 15 minutes before being returned to their home cage.

### Immunohistochemistry and Confocal Microscopy

For all analyses of barrel cortex, mice were perfused transcardially with ∼10 mL of ice-cold PBS followed by ∼10 mL of 4% (weight/volume) paraformaldehyde diluted in PBS. For analyses of SARS-CoV-2 infected brains, animals were not perfused and were postfixed by immersion in 4% PFA for 72 hours prior to cryoprotection and sectioning. All other brains were post-fixed in 4% PFA for a minimum of 4 hours and then transferred to a 30% sucrose solution for a minimum of 24 hours. Tangential sections (flat mounts) of the barrel cortex were obtained by dissecting the cortex from the diencephalon following perfusion and flattening the dissected cortices ventral side down between two cryomolds. The sections were placed in between flat toothpicks laid horizontally in the cryomold, and the second mold was pressed down on top of the toothpicks to maintain uniform thickness. Brains were then flash frozen and sliced on a HM 440E freezing microtome (GMI Instruments) or embedded in OCT following 30% sucrose treatment and frozen at −80°C for a minimum of 1 day and then sectioned on a CryoStar NX70 Cryostat (Thermo Fisher) before being mounted on coverslips. Sections from control and deprived hemispheres of individual mice were mounted on the same slides for all experiments.

### See resource table below for details of all antibodies used

Immunohistochemistry was performed as follows: brain sections were incubated in a blocking solution consisting of 5% normal goat serum (Thermo Fisher) and 0.1% Triton (Sigma-Aldrich) diluted in 1X PBS. Primary antibodies were diluted in 5% normal goat serum in 0.1% Triton and tissue was incubated on a shaker overnight at 4°C. Secondary antibodies were diluted in 5% normal goat serum and tissue was incubated on a shaker for 2 hours at room temperature. Brain sections were mounted on coverslips with ProLong Gold or Glass (Thermo Fisher) for all imaging. For staining with IFITM3 (Thermo Fisher #11714-1-AP), secondary antibody staining was done with goat anti-rabbit IgG, HRP-linked (Cell Signaling Technology #7074) and visualized with TSA Plus Cy3 detection system (Akoya Biosciences #NEL744001KT). For staining with SARS-CoV-2 Spike and N protein antibodies (GeneTex), an additional antigen retrieval step (70°C for 10min in 1X citrate buffer) was performed prior to blocking. For staining with 53BP1 and γH2AX, 1X TBS was used instead of 1X PBS and additional quenching step (50 mM NH4Cl/TBS for 5 min at RT) and permeabilization (1% TX-100/TBS for 20 min on ice) were performed prior to primary antibody incubation. Slides were imaged on an LSM 800 confocal microscope (Zeiss, Zen 2.6 software) using 20x, 40x, and 63x objectives. For analysis of phagocytic cups, slides were imaged on LSM 880 confocal microscope with AiryScan (Zeiss) using a 63x objective.

### Fluorescent In Situ Hybridization (FISH)

FISH experiments were performed using the RNAscope Multiplex Fluorescent Reagent Kit v1 assay for *Htra1* and v2 assay for *Grin1* and *Rbfox3* (ACD Bio) as described by the manufacturer for fixed-frozen tissue, but eliminating the 60°C incubation and post-fixation steps prior to tissue dehydration. Brains were embedded in OCT following 30% sucrose treatment and frozen at −80°C for a minimum of 1 day prior to sectioning. Mouse *Htra1* RNAscope Probe (ACD Bio #423711-C2), Mouse *Grin1* RNAscope Probe (ACD Bio #431611-C1), Mouse *Rbfox3* Probe (ACD Bio #313311-C2) and Mouse *Ifnar1* RNAscope probe (ACD Bio #512971-C2) were used to detect each transcript. For immunohistochemical labeling with antibodies following the RNAscope assay, tissues were incubated with blocking and antibody solutions as described above immediately after RNAscope and washing four times, 5 minutes each. Confocal optical sections were imaged on a Zeiss 700 at 63x magnification through layer IV of flattened *en face* cortical sections of the barrel cortex. The Htra1 channel was thresholded to remove puncta smaller than 0.06 um^2 in area, and the puncta within a 10mm radius of each Aldh1l1-GFP+ astrocyte were counted. Three images each from at least two sections each of three separate mice were counted per condition.

### Flow cytometry

For all flow cytometry experiments, animals were perfused transcardially with ice-cold D-PBS, mounted coronally in ice-cold isolation media (HBSS, 15 mM HEPES, 0.6% glucose, 1 mM EDTA pH 8.0) and 350 micrometer slices were prepared on a vibratome. A stereomicroscope was used to visually identify barrel regions; 1 mm^3^ of tissue was collected per hemisphere. All data analysis was performed using FlowJo^TM^ software.

### MACS Bead Isolation

For sorting microglia only for downstream RNA-sequencing, cells were isolated as described previously^84^. Briefly, barrel cortices dissected as described above were mechanically dissociated using a glass tissue homogenizer in isolation medium (HBSS, 15 mM HEPES, 0.6% glucose, 1 mM EDTA pH 8.0). Cells were filtered and then pelleted at 300 g for 10 minutes at 4°C before being resuspended in 22% Percoll (GE Healthcare) and centrifuged at 900 g for 20 minutes with acceleration set to 4 and deceleration set to 1 in order to remove cellular debris. Pelleted microglia were then resuspended in staining media (PBS, 0.5% BSA, 2 mM EDTA) and incubated with CD11b MACS beads (Miltenyi, 1:50) for 15 minutes at 4°C. The cells were washed with staining buffer, pelleted at 300 g for 5 minutes at 4°C, and reconstituted in 500 uL staining buffer. Microglia were isolated as described in the manual for MACS LS columns and collected in staining buffer without EDTA, pelleted at 300 g for 5 minutes at 4°C, counted on a hemocytometer, and 15,000-20,000 cells were diluted in 30 µL in a BSA-coated plate for 10x sequencing.

### Microglial single cell sequencing (Fig. 1)

Single cells were isolated as described above. Approximately 15,000 cells were loaded into each well of Chromium Chip B, libraries were prepared in-house as described in the 10x Manual, and sequenced on one lane of the NovaSeq S4 at the Chan-Zuckerberg BioHub.

### Single cell data analysis

Sequenced samples were processed using the Cell Ranger 2.1 pipeline (built on the STAR aligner)^85^ and aligned to the GRCm38 (mm10) mouse reference genome. Clustering and differential expression analysis were conducted using Seurat version 3.1.4. Data for figures 1 and 2 (total glial population) and figure 3 (microglia only) were prepared on different versions of the 10x Chromium platform and were therefore analyzed separately as detailed below. Sequencing scripts can be found at https://github.com/lcdorman/IFNresponseCode, and original data can be found on GEO at <u>GSE173173</u>.

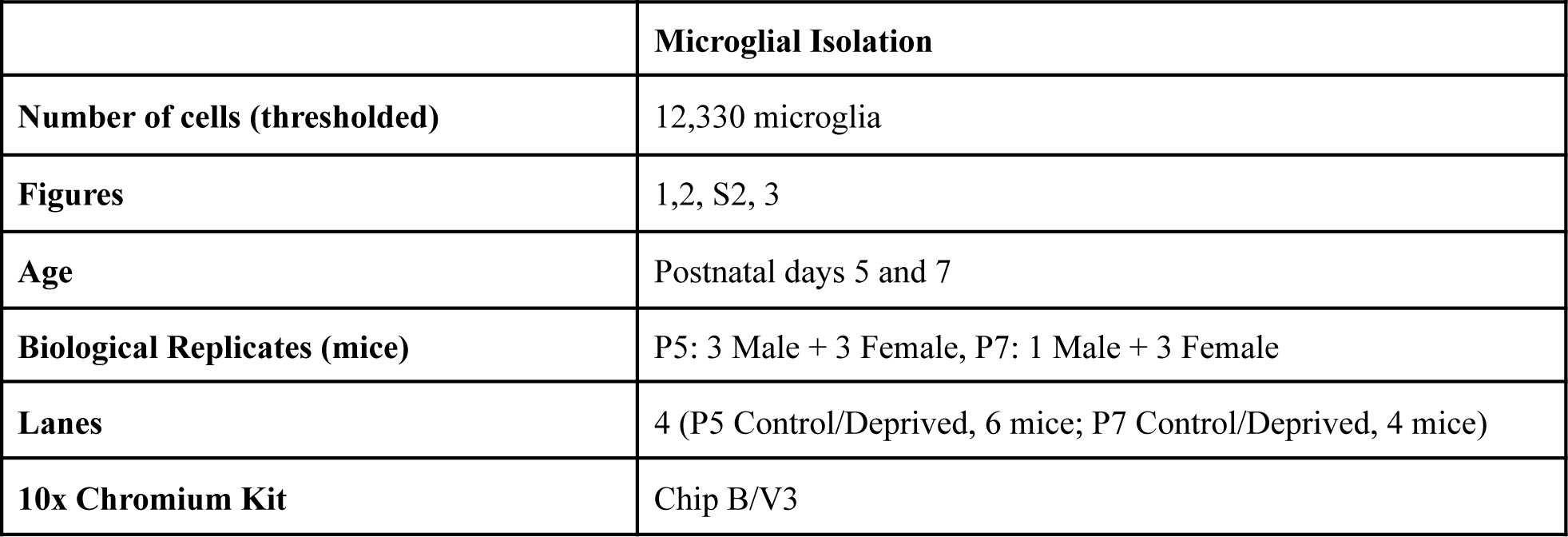

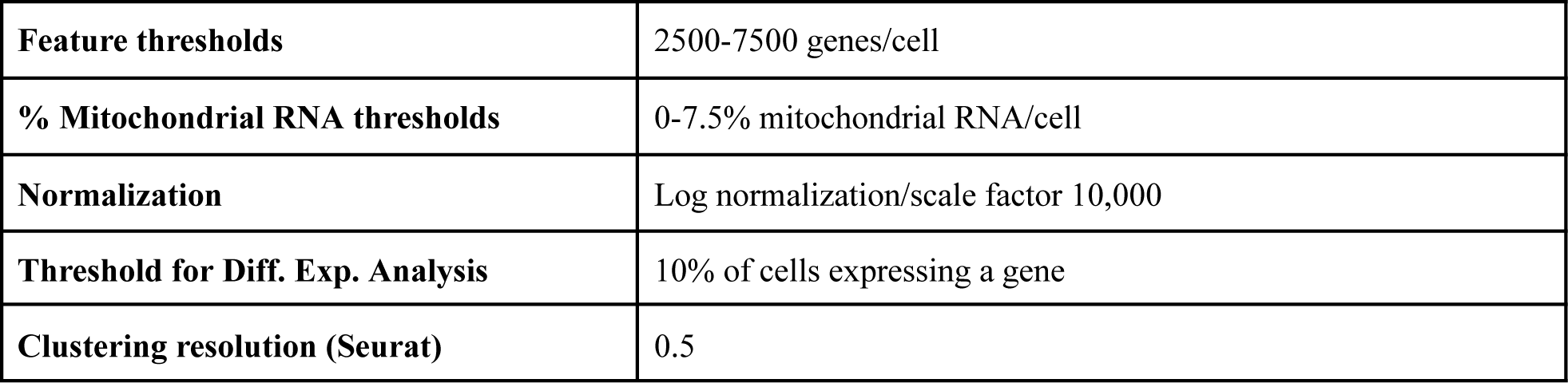

Sorted cells were sequenced using the 10x Chromium kit. Following alignment in Cell Ranger as described above, counts were imported into R and analyzed using the Seurat package^86,87^. Cells outside of the thresholds listed in the table were excluded from downstream analysis. Cells were identified as “female” or “male” based on their expression of the gene *Xist*; any cells expressing at least one count of *Xist* were labeled female, while all others were labeled male. Counts were then normalized, regressing out percent mitochondrial RNA and total counts per cell. The top 6000 most variable genes were used to calculate 50 principal components, and the top 30 PCs were used for nearest neighbor, UMAP, and cluster calculations with the resolutions shown in the table. Individual cell types were identified through calculation of marker genes using the Wilcox test for genes expressed in at least 50% of cells in the cluster and a natural log fold change of 1.2 or greater and adjusted p value less than 0.001.

Microglial and macrophage clusters were isolated based on expression of Cx3cr1, Fcrls, P2y12, and low expression of non-microglial genes as shown in the heatmap. The filtered cells were re-normalized and analyzed with the Variance Partition package in R to determine the top genes determining sex, the first 8 of which were excluded from downstream analysis. The top 6000 most variable genes were used to recalculate PCs, UMAP, and clusters. Clusters were determined using a resolution of 0.5, and the two sets of most closely related clusters (0/5 and 6/7) were combined due to low numbers of unique differentially expressed genes (log fold change >0.15, adjusted p-value < 10^-5^). Clusters were combined either due to relatively low numbers of uniquely upregulated genes (<30 between clusters 6/7, 34 between clusters 0 and 5). The resultant and remaining clusters had equal or more unique DEGs and importantly were not more closely related to any other clusters with few unique DEGs. Differential gene expression between clusters were calculated using the MAST test in Seurat. The heatmaps shown only include genes expressed by at least 50% of the cells in that cluster and with an adjusted p-value below 10^-25^, sorted by highest log fold change. GO analysis was conducted using the Metascape webpage (www.metascape.org)81. Volcano plots were generated using the EnhancedVolcano package in R, with gene labels chosen from the top differentially expressed genes. Cutoffs were set at natural log fold change greater than 0.2 (22% increase) and adjusted p-value smaller than 10^-25^. Gene signature enrichment scores (Figure S2K) were calculated using Seurat’s PercentageFeatureSet function.

#### Cell cycle phase assignment

Cells were assigned to S phase, G1 phase, or G2/M phase (not distinguished) using a previously published dataset^82^ and the CellCycleScoring function in Seurat.

#### RNA Velocity analysis

Spliced and unspliced transcript counts were calculated using Velocyto 0.17 and the velocyto run10x command with default settings. UMAP cell embeddings and annotation were exported from Seurat and used to plot all data shown. ScVelo 2.0.0 was run in Python and trajectories were calculated for all cells, then for each sample individually^54,88^.

*Bar plot creation*: Bar plots were created using ggplot2 in R. Coding details are available on github at (https://github.com/lcdorman/IFNresponseCode/blob/main/Code%20for%20paper/P5_P7%20Microglia/D_BarPlots.Rmd). A table was made of cells per cluster per sample. Cell numbers were normalized by sample by dividing each entry by the total number of cells for that sample and multiplying by 2,000. Percents per cluster were then calculated by dividing the normalized cell numbers by the total number of cells in that cluster and multiplying by 100. Statistics were calculated individually for each cluster using a Chi-Square test on the raw cell numbers per cluster and sample. Plots with multiple bars had an additional Bonferroni correction applied, which multiplies the p-value by the number of comparisons.

### qPCR

To extract RNA from cells isolated by FACS, freshly sorted cells were pelleted at 500 g for 10 minutes at 4° and then resuspended in RLT Plus buffer (Qiagen). Cells were vortexed and frozen for at least one day at −80° before being thawed on ice and processed for RNA using an RNeasy Mini Kit (Qiagen). Purified mRNA was converted to cDNA with the High Capacity cDNA Reverse Transcription kit (Life Technologies) and amplified using either the Fast SYBR Green Master Mix (Thermo Fisher) or TaqMan Gene Expression Master Mix (Thermo Fisher) and a 7900HT Fast Real-Time PCR System (Applied Biosystems).

### SARS-CoV-2 virus propagation and plaque assay

All SARS-CoV-2 cell culture and animals works were performed in the Biosafety level 3(BSL3). African green monkey kidney Vero-E6 cell line (ATCC#1586) and Calu-3 cells (ATCC# HTB-55) was obtained from American Type Culture Collection and maintained in Minimum Essential Medium (MEM, Gibco Invitrogen) supplemented with 10% fetal bovine serum (FBS, Gibco Invitrogen), 1% Penicillin-Streptomycin-Glutamine (Gibco Invitrogen) at 37 °C in a humidified 5% CO2 incubator. A clinical isolate of SARS-CoV-2 from a UCSF patient was propagated in Vero E6 cells and Calu-3 cells. 80% Confluent monolayers of Vero E6 cells grown in 6-well plates were incubated with the serial dilutions of virus samples (250 ul/well) at 37 °C for 1 hour. Next, the cells were overlayed with 1% agarose (Invitrogen) prepared with MEM supplemented containing 2% fetal bovine serum(sigma), 1x penicillin/streptomycin/glutamine (100xPSG, Gibco). Three days later, the cells were fixed with 4% formaldehyde (PFA) for 2 hours, the overlay was discarded and samples were stained with crystal violet dye.

### SARS-CoV-2 infection

5-6 weeks old hemizygous K18-hACE2 mice (The Jackson laboratory, https://www.jax.org/strain/034860, stock number: 034860, B6.Cg-Tg(K18-ACE2)2Prlmn/J) were anesthetized with isofluorane and inoculated with 6x10^4^ PFU of SARS-CoV-2 intranasally in a BSL3 facility. The mice were sacrificed at 3 and 6 days post-infection, and the brain was removed and fixed in 4% PFA for 72 hours before being sunk in 30% sucrose and embedded in OCT. 40 µm sections were cut on a cryostat and stained as described above with 2 minute antigen retrieval at 95 degrees. Antibodies used include the following: rabbit anti-SARS-CoV-2 Nucleocapsid(N) (GeneTex, GTX135361), mouse anti-SARS-CoV-2-spike(S), (GeneTex, GTX632604).

### Microglia CD68 volume

Z-stacks were collected on an LSM 880 confocal microscope with AiryScan (Zeiss) on Superresolution mode and a 63x objective (NA 1.4). Laser power and gain were consistent across each image. AiryScan processing was performed in Zen software (Zeiss) at a setting of 6 (‘‘optimal’’ setting). Images were analyzed using Imaris software (Bitplane) by creating a 3D surface rendering of individual microglia, thresholded to ensure microglia processes were accurately reconstructed, and maintained consistent thereafter. Microglia rendering was used to mask and render the CD68 channel within each microglia. CD68 volume per microglia was then calculated as the total volume of masked CD68 volume within the masked GFP volume.

### Microglia phagocytic compartment analyses

Phagocytic compartments, including phagocytic cups, phagosomes, and phagolysosomes, were identified as DAPI-enveloping structures that are distinct from the microglia nuclei. Unlike the microglia nuclear compartment, phagosomes lacked staining for IBA1 or GFP. Phagocytic cups lacked CD68 and only partially enveloped a DAPI+ and non-pyknotic cell. Microglia with bubble morphology were identified by enlarged and rounded phagosomes or phagolysosomes that were larger than the microglia nucleus. These phagocytic compartments contained DAPI+ nuclear material undergoing pyknosis or karyorrhexis, which were very sparsely distributed within the phagosome. In contrast, non-bubble phagosomes tightly enveloped engulfed nuclear material such that the DAPI signal saturated the phagosome area.

Z-stacks were acquired using an LSM 880 confocal microscope with AiryScan (Zeiss) on Superresolution mode and a 63x objective (NA 1.4, 0.04 _mm_ pixel size, 16-bit depth) and processed as described above. Phagocytic compartments were analyzed in ImageJ and ROI’s were drawn around phagosomes using the Versatile Wand Tool plugin. The “tolerance” setting was manually adjusted to envelop the microglia phagocytic compartment surface and “connectedness” was set to 8-connected. An optical section was selected from the z-stack that represented the center of the compartment and mean intensity, integrated density, and area were then recorded for each phagocytic compartment. For analysis of phagocytic compartments in Sars-CoV-2 infected mice, images were first binned by average IFITM3 MFI per microglia. The maximum microglial IFITM3 mean fluorescence intensity per image across all samples was called 100%. The lower bin represents images with 0-50% of the maximum expression, and the higher bin represents images with 50-100% of the maximum expression. Each microglia was then manually scored for presence or absence of a phagocytic compartment as described above.

### Interferon β injection

A solution of 10 ng interferon-β (IFN-β, R&D Systems, 8234-MB-010) diluted in PBS (2μl) was slowly delivered by intracerebroventricular injection into P4 C57/Bl6J mice (coordinates: x = 1, y=1.8, z = −2, from lambda) using a pulled and beveled glass pipette with 50 μm outer diameter. Control mice were injected with an equal volume of PBS. The mice were perfused 24 hours later with ice-cold PBS followed by 4% paraformaldehyde, post-fixed in 4% PFA overnight at 4°C, cryoprotected and sectioned on a cryostat at 50μm (floating sections, used for IFITM3 quantification) and 20μm (slide-mounted sections, used for 53BP1 foci quantification). Images were acquired using the 40x objective of an LSM 800 (Zeiss) and 53BP1 foci+ neurons were manually counted using the Cell Counter tool in FIJI. 53BP1 foci+ neurons as a percentage of 53BP1I+ nuclei was quantified in L2/3 within the region of IFITM3+ expression, which extended approximately 300μm from the injection site over primary and secondary motor cortex.

### Poly(I:C) microinjection into zebrafish larvae

For poly(I:C) injection, 7 dpf *Tg(mpeg:EGFP-CAAX);Tg(NBT:dsRed)* zebrafish larvae were anesthetized with 0.2 mg/ml of tricaine in embryo medium. Larvae were injected with 2 nl of 1 mg/ml poly(I:C) (Invitrogen) or PBS as a vehicle control into the optic tectal ventricle by microinjection. Zebrafish larvae were transferred to fresh embryo medium after injection for recovery and imaged at 4 hours post injection. We carefully monitored the larvae after injection to confirm that they recovered from anesthesia and did not exhibit abnormal swimming behaviors. Only healthy post-injection larvae were used for live imaging.

### Zebrafish Live imaging

For live imaging, *Tg(mpeg:EGFP-CAAX);Tg(NBT:dsRed)* zebrafish larvae were anesthetized with 0.2 mg/ml of tricaine in embryo medium and mounted in 1.2% low-melting agarose gel on a glass bottom 35-mm dish (MatTek) and covered with embryo water containing 0.2 mg/ml tricaine. Time-lapse image was performed on a Nikon CSU-W1 spinning disk/high speed widefield microscope. We took time-lapse images from the optic tectum collecting 40-60 mm z-stacks (step size: 0.5 mm) at 5 min intervals for 30 min-1 hr. The images were processed by ImageJ software. For analysis, we calculated the number of dsRed(+) neurons in microglia for the first 30 min of each video. For the analysis of process motility, we carefully observed each microglia process for the first 30 min of each video and classified all processes into three categories: 1) ‘ramified’ were defined as processes with or without movement that have no phagocytic cup formation, 2) ‘cup formation’ denoted processes with or without movement that contained or formed phagocytic cups, and 3) ‘soma retraction’ - processes that enclosed dsRed+ cells that were subsequently trafficked toward the microglial soma). Statistical analyses were performed by Graphpad Prism software.

## −QUANTIFICATION AND STATISTICAL ANALYSIS

Graphpad Prism 9.3.1 was used for most statistical analyses of imaging data, and the Seurat package V3 in R was used for statistical analysis of single cell data. Statistical tests are as described in text and figure legends. Violin plots were used for data with n > 20 to better visualize the distribution of individual data points. Single cell RNA-sequencing data was analyzed in R as described in the methods section above. Categorical data shown in bar plots (cluster-specific differences in cell numbers) were analyzed in R using a Chi-Square test on each cluster with Bonferroni’s correction for multiple comparisons.

## RESOURCE TABLE

| REAGENT or RESOURCE<br>SOURCE<br>IDENTIFIER |
| --- |
| **Antibodies-immunostaining** |
| rabbit anti-dsRed 1:1000 (Clontech, 632496) |
| Rat anti-mCherry 1:1000 (Invitrogen, M11217) |
| chicken anti-GFP 1:1000 (Aves Labs, 1020) |
| chicken anti-NeuN1:400 (Millipore, ABN91) |
| rabbit anti-Iba1 1:1000 (Wako Chemicals, 019-19741) |
| Mouse anti-Iba1 1:1000 (Wako Chemicals, 016-26721) |
| Guinea pig anti-Iba1 1:2000 (Synaptic Systems, 234-004) |
| Rat anti-P2ry12 1:100 (Biolegend, 848001) |
| rabbit anti-P2y12 1:500 (David Julius Lab) |
| rat anti-GFAP 1:1000 (Thermo Fisher, 13-0300) |
| Rabbit anti-IFITM3 1:1:1000 (Thermo Fisher, 11714-1-AP) |
| guinea pig anti-VGLUT2 1:2000 (Synaptic Systems, 135404) |
| rat anti-CD68 1:500 (BioRad, MCA1957GA) |
| Goat anti-rabbit IgG, HRP-linked (Cell Signaling Technology, 7074P2) |
| Hoechst 33342 solution 1:5000 (Thermo Fisher, 62249) was used for nuclear labeling |
| rabbit anti-SARS-CoV-2-N protein 1:1000 (GeneTex, GTX135361) |
| mouse anti-SARS-CoV-2-Spike 1:1000 (GeneTex, GTX632604) |
| Rat anti-Bst2/Tetherin (R&D Systems, MAB8660-SP) |
| Mouse anti- -H2AX (EMD Millipore, 05-636) |
| Rabbit anti-53BP1 (Novus Biologicals, NB100-304) |
| Rabbit anti-cleaved caspase 3 1:1000 (BD Biosciences, 559565) |

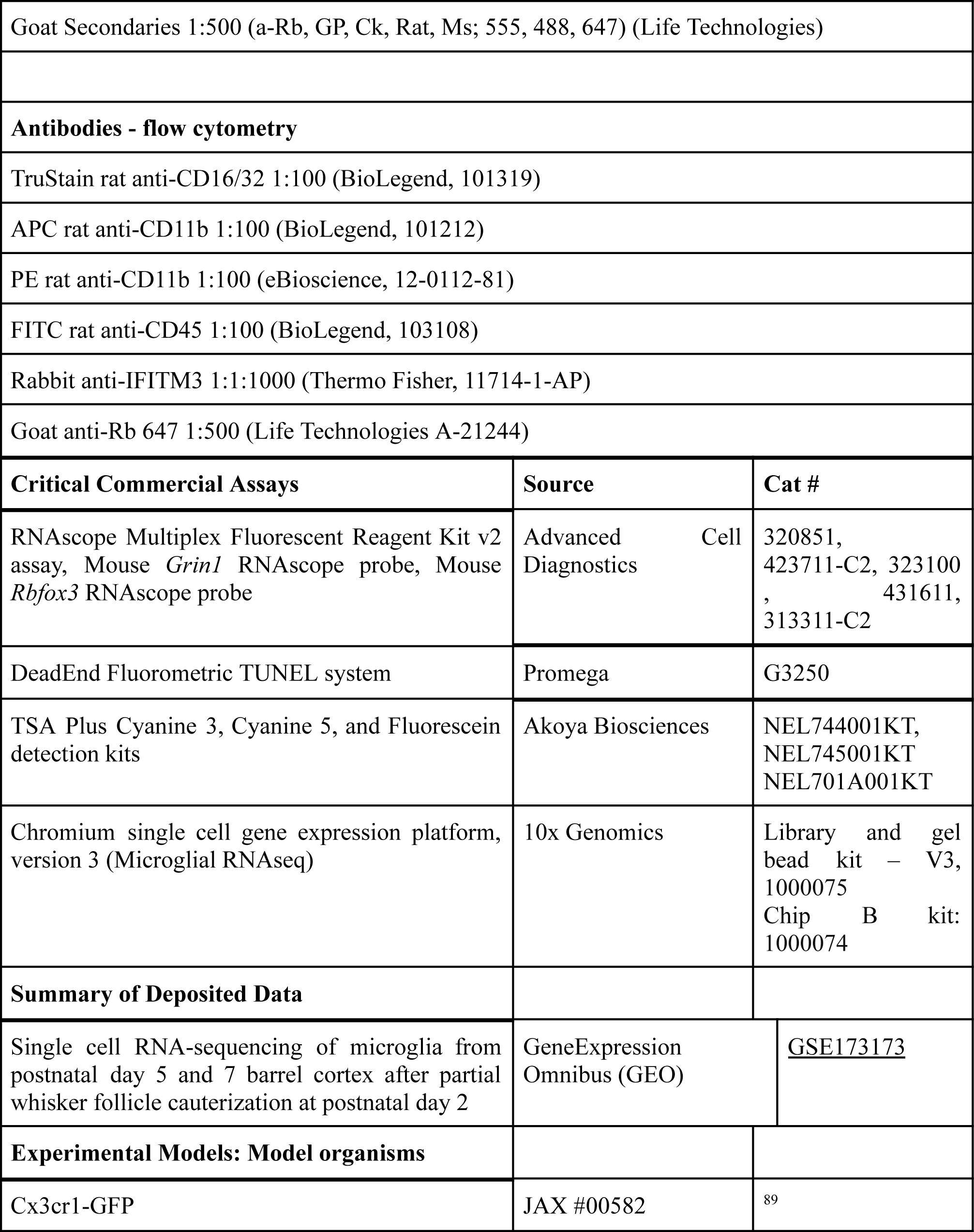

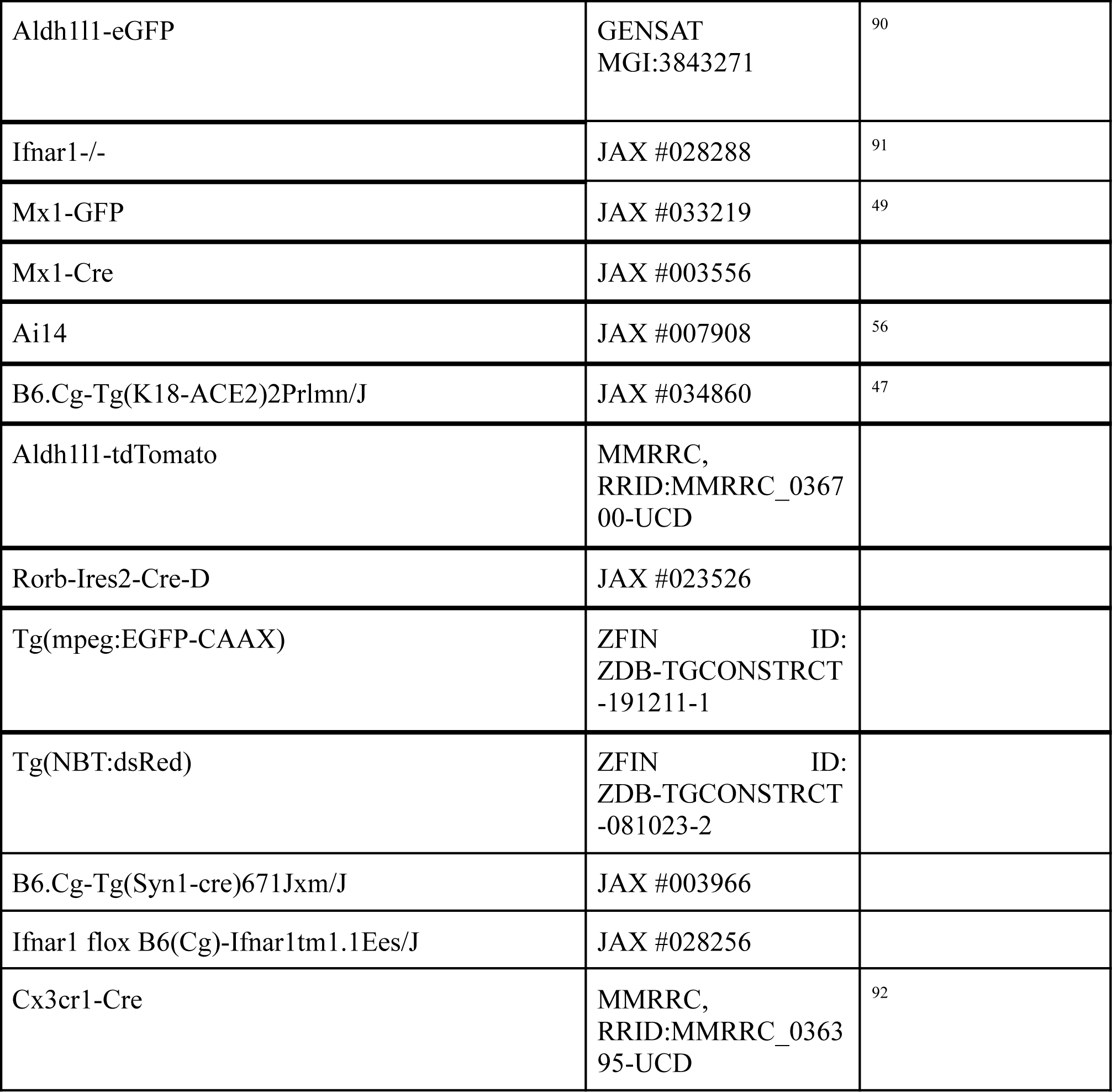

## BIBLIOGRAPHY

1. Forrest, M. P., Parnell, E. & Penzes, P. Dendritic structural plasticity and neuropsychiatric disease. Nat. Rev. Neurosci. 19, 215–234 (2018).

2. Bitzenhofer, S. H., Pöpplau, J. A., Chini, M., Marquardt, A. & Hanganu-Opatz, I. L. A transient developmental increase in prefrontal activity alters network maturation and causes cognitive dysfunction in adult mice. Neuron 109, 1350–1364.e6 (2021).

3. Medendorp, W. E. et al. Selective postnatal excitation of neocortical pyramidal neurons results in distinctive behavioral and circuit deficits in adulthood. iScience 24, 102157 (2021).

4. Arguello, P. A. & Gogos, J. A. Genetic and cognitive windows into circuit mechanisms of psychiatric disease. Trends Neurosci. 35, 3–13 (2012).

5. Vainchtein, I. D. & Molofsky, A. V. Astrocytes and Microglia: In Sickness and in Health. Trends Neurosci. 43, 144–154 (2020).

6. Allen, N. J. & Lyons, D. A. Glia as architects of central nervous system formation and function. Science 362, 181–185 (2018).

7. Bennett, F. C. & Molofsky, A. V. The immune system and psychiatric disease: a basic science perspective. Clin. Exp. Immunol. 197, 294–307 (2019).

8. Prinz, M., Jung, S. & Priller, J. Microglia Biology: One Century of Evolving Concepts. Cell 179, 292–311 (2019).

9. Frost, J. L. & Schafer, D. P. Microglia: Architects of the Developing Nervous System. Trends Cell Biol. 26, 587–597 (2016).

10. Srinivasan, K. et al. Untangling the brain’s neuroinflammatory and neurodegenerative transcriptional responses. Nature Communications vol. 7 Preprint at https://doi.org/10.1038/ncomms11295 (2016).

11. McNab, F., Mayer-Barber, K., Sher, A., Wack, A. & O’Garra, A. Type I interferons in infectious disease. Nat. Rev. Immunol. 15, 87–103 (2015).

12. Gosselin, D. et al. An environment-dependent transcriptional network specifies human microglia identity. Science 356, (2017).

13. Badimon, A. et al. Negative feedback control of neuronal activity by microglia. Nature 586, 417–423 (2020).

14. Van der Loos, H. & Woolsey, T. A. Somatosensory cortex: structural alterations following early injury to sense organs. Science 179, 395–398 (1973).

15. Erzurumlu, R. S. & Gaspar, P. Development and critical period plasticity of the barrel cortex. Eur. J. Neurosci. 35, 1540–1553 (2012).

16. Sehara, K. & Kawasaki, H. Neuronal circuits with whisker-related patterns. Mol. Neurobiol. 43, 155–162 (2011).

17. Woolsey, T. A. & Wann, J. R. Areal changes in mouse cortical barrels following vibrissal damage at different postnatal ages. J. Comp. Neurol. 170, 53–66 (1976).

18. Jeanmonod, D., Rice, F. L. & Van der Loos, H. Mouse somatosensory cortex: alterations in the barrelfield following receptor injury at different early postnatal ages. Neuroscience 6, 1503–1535 (1981).

19. Sehara, K. et al. Whisker-related axonal patterns and plasticity of layer 2/3 neurons in the mouse barrel cortex. J. Neurosci. 30, 3082–3092 (2010).

20. Hoshiko, M., Arnoux, I., Avignone, E., Yamamoto, N. & Audinat, E. Deficiency of the microglial receptor CX3CR1 impairs postnatal functional development of thalamocortical synapses in the barrel cortex. J. Neurosci. 32, 15106–15111 (2012).

21. Gunner, G. et al. Sensory lesioning induces microglial synapse elimination via ADAM10 and fractalkine signaling. Nat. Neurosci. 22, 1075–1088 (2019).

22. Pollina, E. A. et al. A NPAS4-NuA4 complex couples synaptic activity to DNA repair. Nature 614, 732–741 (2023).

23. Madabhushi, R. et al. Activity-Induced DNA Breaks Govern the Expression of Neuronal Early-Response Genes. Cell 161, 1592–1605 (2015).

24. Suberbielle, E. et al. Physiologic brain activity causes DNA double-strand breaks in neurons, with exacerbation by amyloid-β. Nat. Neurosci. 16, 613–621 (2013).

25. Marek, R., Caruso, M., Rostami, A., Grinspan, J. B. & Das Sarma, J. Magnetic cell sorting: a fast and effective method of concurrent isolation of high purity viable astrocytes and microglia from neonatal mouse brain tissue. J. Neurosci. Methods 175, 108–118 (2008).

26. Bennett, F. C. et al. A Combination of Ontogeny and CNS Environment Establishes Microglial Identity. Neuron 98, 1170–1183.e8 (2018).

27. Hammond, T. R. et al. Single-Cell RNA Sequencing of Microglia throughout the Mouse Lifespan and in the Injured Brain Reveals Complex Cell-State Changes. Immunity 50, 253–271.e6 (2019).

28. Li, Q. et al. Developmental Heterogeneity of Microglia and Brain Myeloid Cells Revealed by Deep Single-Cell RNA Sequencing. Neuron 101, 207–223.e10 (2019).

29. Keren-Shaul, H. et al. A Unique Microglia Type Associated with Restricting Development of Alzheimer’s Disease. Cell 169, 1276–1290.e17 (2017).

30. Friedman, B. A. et al. Diverse Brain Myeloid Expression Profiles Reveal Distinct Microglial Activation States and Aspects of Alzheimer’s Disease Not Evident in Mouse Models. Cell Rep. 22, 832–847 (2018).

31. Sala Frigerio, C., et al. The Major Risk Factors for Alzheimer’s Disease: Age, Sex, and Genes Modulate the Microglia Response to Aβ Plaques. Cell Rep. 27, 1293–1306.e6 (2019).

32. Zhan, L. et al. A MAC2-positive progenitor-like microglial population is resistant to CSF1R inhibition in adult mouse brain. Elife 9, (2020).

33. Ochocka, N. et al. Single-cell RNA sequencing reveals functional heterogeneity of glioma-associated brain macrophages. Nat. Commun. 12, 1151 (2021).

34. Masuda, T. et al. Spatial and temporal heterogeneity of mouse and human microglia at single-cell resolution. Nature 566, 388–392 (2019).

35. Shi, L. et al. Treg cell-derived osteopontin promotes microglia-mediated white matter repair after ischemic stroke. Immunity 54, 1527–1542.e8 (2021).

36. Lee, S.-H. et al. Trem2 restrains the enhancement of tau accumulation and neurodegeneration by β-amyloid pathology. Neuron 109, 1283–1301.e6 (2021).

37. Syage, A. R. et al. Single-Cell RNA Sequencing Reveals the Diversity of the Immunological Landscape following Central Nervous System Infection by a Murine Coronavirus. J. Virol. 94, (2020).

38. Hur, J.-Y. et al. The innate immunity protein IFITM3 modulates γ-secretase in Alzheimer’s disease. Nature 586, 735–740 (2020).

39. Bailey, C. C., Zhong, G., Huang, I.-C. & Farzan, M. IFITM-Family Proteins: The Cell’s First Line of Antiviral Defense. Annu Rev Virol 1, 261–283 (2014).

40. Brass, A. L. et al. The IFITM proteins mediate cellular resistance to influenza A H1N1 virus, West Nile virus, and dengue virus. Cell 139, 1243–1254 (2009).

41. Zhang, Y. et al. Interferon-Induced Transmembrane Protein 3 Genetic Variant rs12252-C Associated With Disease Severity in Coronavirus Disease 2019. J. Infect. Dis. 222, 34–37 (2020).

42. Shi, G., et al. Opposing activities of IFITM proteins in SARS-CoV-2 infection. The EMBO Journal vol. 40 Preprint at https://doi.org/10.15252/embj.2020106501 (2021).

43. Liotta, E. M. et al. Frequent neurologic manifestations and encephalopathy-associated morbidity in Covid-19 patients. Ann Clin Transl Neurol 7, 2221–2230 (2020).

44. Taquet, M., Luciano, S., Geddes, J. R. & Harrison, P. J. Bidirectional associations between COVID-19 and psychiatric disorder: retrospective cohort studies of 62 354 COVID-19 cases in the USA. Lancet Psychiatry 8, 130–140 (2021).

45. Moreau, G. B. et al. Evaluation of K18-hACE2 Mice as a Model of SARS-CoV-2 Infection. The American Journal of Tropical Medicine and Hygiene vol. 103 1215–1219 Preprint at https://doi.org/10.4269/ajtmh.20-0762 (2020).

46. Winkler, E. S. et al. SARS-CoV-2 infection of human ACE2-transgenic mice causes severe lung inflammation and impaired function. Nat. Immunol. 21, 1327–1335 (2020).

47. McCray, P. B., Jr et al. Lethal infection of K18-hACE2 mice infected with severe acute respiratory syndrome coronavirus. J. Virol. 81, 813–821 (2007).

48. Kumari, P. et al. Neuroinvasion and Encephalitis Following Intranasal Inoculation of SARS-CoV-2 in K18-hACE2 Mice. Viruses 13, (2021).

49. Uccellini, M. B. & García-Sastre, A. ISRE-Reporter Mouse Reveals High Basal and Induced Type I IFN Responses in Inflammatory Monocytes. Cell Rep. 25, 2784–2796.e3 (2018).

50. Liberatore, R. A. & Bieniasz, P. D. Tetherin is a key effector of the antiretroviral activity of type I interferon in vitro and in vivo. Proc. Natl. Acad. Sci. U. S. A. 108, 18097–18101 (2011).

51. Chistiakov, D. A., Killingsworth, M. C., Myasoedova, V. A., Orekhov, A. N. & Bobryshev, Y. V. CD68/macrosialin: not just a histochemical marker. Lab. Invest. 97, 4–13 (2017).

52. Uribe-Querol, E. & Rosales, C. Phagocytosis: Our Current Understanding of a Universal Biological Process. Front. Immunol. 11, 1066 (2020).

53. Lancaster, C. E. et al. Phagosome resolution regenerates lysosomes and maintains the degradative capacity in phagocytes. J. Cell Biol. 220, (2021).

54. Bergen, V., Lange, M., Peidli, S., Wolf, F. A. & Theis, F. J. Generalizing RNA velocity to transient cell states through dynamical modeling. Nat. Biotechnol. 38, 1408–1414 (2020).

55. Harris, J. A. et al. Anatomical characterization of Cre driver mice for neural circuit mapping and manipulation. Front. Neural Circuits 8, 76 (2014).

56. Madisen, L. et al. A robust and high-throughput Cre reporting and characterization system for the whole mouse brain. Nat. Neurosci. 13, 133–140 (2010).

57. Mah, L.-J., El-Osta, A. & Karagiannis, T. C. gammaH2AX: a sensitive molecular marker of DNA damage and repair. Leukemia 24, 679–686 (2010).

58. Villani, A. et al. Clearance by Microglia Depends on Packaging of Phagosomes into a Unique Cellular Compartment. Dev. Cell 49, 77–88.e7 (2019).

59. Secombes, C. J. & Zou, J. Evolution of Interferons and Interferon Receptors. Front. Immunol. 8, 209 (2017).

60. Boudinot, P., Langevin, C., Secombes, C. J. & Levraud, J.-P. The Peculiar Characteristics of Fish Type I Interferons. Viruses 8, (2016).

61. Silva, N. J., Dorman, L. C., Vainchtein, I. D., Horneck, N. C. & Molofsky, A. V. In situ and transcriptomic identification of microglia in synapse-rich regions of the developing zebrafish brain. Nature Communications vol. 12 Preprint at https://doi.org/10.1038/s41467-021-26206-x (2021).

62. Peri, F. & Nüsslein-Volhard, C. Live Imaging of Neuronal Degradation by Microglia Reveals a Role for v0-ATPase a1 in Phagosomal Fusion In Vivo. Cell vol. 133 916–927 Preprint at https://doi.org/10.1016/j.cell.2008.04.037 (2008).

63. Wong, F. K. & Marín, O. Developmental Cell Death in the Cerebral Cortex. Annu. Rev. Cell Dev. Biol. 35, 523–542 (2019).

64. Harb, K. et al. Area-specific development of distinct projection neuron subclasses is regulated by postnatal epigenetic modifications. Elife 5, e09531 (2016).

65. McNamara, K. C. S., Lisembee, A. M. & Lifshitz, J. The whisker nuisance task identifies a late-onset, persistent sensory sensitivity in diffuse brain-injured rats. J. Neurotrauma 27, 695–706 (2010).

66. Balasco, L., Chelini, G., Bozzi, Y. & Provenzano, G. Whisker Nuisance Test: A Valuable Tool to Assess Tactile Hypersensitivity in Mice. Bio Protoc 9, e3331 (2019).

67. Chelini, G. et al. Aberrant Somatosensory Processing and Connectivity in Mice Lacking*Engrailed-2*. The Journal of Neuroscience vol. 39 1525–1538 Preprint at https://doi.org/10.1523/jneurosci.0612-18.2018 (2019).

68. Ivashkiv, L. B. & Donlin, L. T. Regulation of type I interferon responses. Nat. Rev. Immunol. 14, 36–49 (2014).

69. Brendecke, S. M. & Prinz, M. How type I interferons shape myeloid cell function in CNS autoimmunity. J. Leukoc. Biol. 92, 479–488 (2012).

70. Kumaran Satyanarayanan, S., et al. IFN-β is a macrophage-derived effector cytokine facilitating the resolution of bacterial inflammation. Nat. Commun. 10, 3471 (2019).

71. Li, X. et al. Viral DNA Binding to NLRC3, an Inhibitory Nucleic Acid Sensor, Unleashes STING, a Cyclic Dinucleotide Receptor that Activates Type I Interferon. Immunity 50, 591–599.e6 (2019).

72. Takaoka, A. & Yamada, T. Regulation of signaling mediated by nucleic acid sensors for innate interferon-mediated responses during viral infection. Int. Immunol. 31, 477–488 (2019).

73. Boada-Romero, E., Martinez, J., Heckmann, B. L. & Green, D. R. The clearance of dead cells by efferocytosis. Nat. Rev. Mol. Cell Biol. 21, 398–414 (2020).

74. Chen, Q., Sun, L. & Chen, Z. J. Regulation and function of the cGAS-STING pathway of cytosolic DNA sensing. Nat. Immunol. 17, 1142–1149 (2016).

75. Knuesel, I. et al. Maternal immune activation and abnormal brain development across CNS disorders. Nat. Rev. Neurol. 10, 643–660 (2014).

76. Estes, M. L. & McAllister, A. K. Immune mediators in the brain and peripheral tissues in autism spectrum disorder. Nat. Rev. Neurosci. 16, 469–486 (2015).

77. Website. American Psychiatric Association. (2013). Diagnostic and statistical manual of mental disorders (5th ed.). https://doi.org/10.1176/appi.books.9780890425596.

78. Mammen, M. A. et al. INFANT AVOIDANCE DURING A TACTILE TASK PREDICTS AUTISM SPECTRUM BEHAVIORS IN TODDLERHOOD. Infant Ment. Health J. 36, 575–587 (2015).

79. Orefice, L. L. et al. Peripheral Mechanosensory Neuron Dysfunction Underlies Tactile and Behavioral Deficits in Mouse Models of ASDs. Cell 166, 299–313 (2016).

80. Malkova, N. V., Yu, C. Z., Hsiao, E. Y., Moore, M. J. & Patterson, P. H. Maternal immune activation yields offspring displaying mouse versions of the three core symptoms of autism. Brain Behav. Immun. 26, 607–616 (2012).

81. Zhou, Y. et al. Metascape provides a biologist-oriented resource for the analysis of systems-level datasets. Nat. Commun. 10, 1523 (2019).

82. Kowalczyk, M. S. et al. Single-cell RNA-seq reveals changes in cell cycle and differentiation programs upon aging of hematopoietic stem cells. Genome Res. 25, 1860–1872 (2015).

83. Zhang, Y. et al. An RNA-sequencing transcriptome and splicing database of glia, neurons, and vascular cells of the cerebral cortex. J. Neurosci. 34, 11929–11947 (2014).

84. Galatro, T. F., Vainchtein, I. D., Brouwer, N., Boddeke, E. W. G. M. & Eggen, B. J. L. Isolation of Microglia and Immune Infiltrates from Mouse and Primate Central Nervous System. Methods Mol. Biol. 1559, 333–342 (2017).

85. Butler, A., Hoffman, P., Smibert, P., Papalexi, E. & Satija, R. Integrating single-cell transcriptomic data across different conditions, technologies, and species. Nat. Biotechnol. 36, 411–420 (2018).

86. Macosko, E. Z. et al. Highly Parallel Genome-wide Expression Profiling of Individual Cells Using Nanoliter Droplets. Cell 161, 1202–1214 (2015).

87. Hoffman, G. E. & Schadt, E. E. variancePartition: interpreting drivers of variation in complex gene expression studies. BMC Bioinformatics 17, 483 (2016).

88. La Manno, G. et al. RNA velocity of single cells. Nature 560, 494–498 (2018).

89. Jung, S. et al. Analysis of fractalkine receptor CX(3)CR1 function by targeted deletion and green fluorescent protein reporter gene insertion. Mol. Cell. Biol. 20, 4106–4114 (2000).

90. Gong, S. et al. A gene expression atlas of the central nervous system based on bacterial artificial chromosomes. Nature 425, 917–925 (2003).

91. Prigge, J. R. et al. Type I IFNs Act upon Hematopoietic Progenitors To Protect and Maintain Hematopoiesis during Pneumocystis Lung Infection in Mice. J. Immunol. 195, 5347–5357 (2015).

92. Zhao, X.-F. et al. Targeting Microglia Using Cx3cr1-Cre Lines: Revisiting the Specificity. eNeuro 6, (2019).

